# YY1 phosphorylation links non-apoptotic caspase-8 signaling to MASH fibrosis

**DOI:** 10.64898/2026.10.05.756806

**Authors:** Xiaobo Wang, Eui Jung Jung, Lanuza A.P. Faccioli, Dwayne G. Stupack, Alejandro Soto-Gutierrez, Ira Tabas

## Abstract

**Background & Aims:** Metabolic dysfunction-associated steatohepatitis (MASH) is a leading cause of chronic liver disease, and liver fibrosis is the principal determinant of disease progression and clinical outcome. We recently showed that non-apoptotic caspase-8 in hepatocytes is upregulated in MASH and leads to the induction and secretion of meteorin, which promotes hepatic stellate cell (HSC) activation and liver fibrosis. We address here the major mechanistic question arising from this study, namely how non-apoptotic caspase-8 induces pro-fibrotic meteorin in hepatocytes in MASH.

**Methods:** Mechanistic studies were performed using AML12 hepatocytes; primary mouse and human hepatocytes; and diet-induced mouse models of MASH with hepatocyte-specific caspase-8 deletion. YY1 post-translational modifications were identified by LC–MS/MS and investigated using phosphorylation-defective and phosphomimetic YY1 mutants. Genetic and biochemical approaches were used to interrogate the caspase-8 DED–SRC–GSK3β pathway, and its relevance was assessed in human MASH liver specimens.

**Results:** In vitro mechanistic stduies showed that the amino-terminal death effector domain (DED) of caspase-8 activated a SRC–GSK3β signaling cascade that promoted phosphorylation of YY1 at Ser247, resulting in YY1 nuclear localization and *Metrn* induction. Disruption of the conserved DED RXDLL motif impaired SRC activation, YY1 phosphorylation, and *Metrn* expression. Hepatocyte YY1 overexpression increased *Metrn* expression and fibrosis in wild-type MASH mice but not in hepatocyte-caspase-8-deficient mice. In contrast, phosphomimetic YY1-S247D restored these endpoints even when hepatocyte caspase-8 was lacking. Studies in primary mouse and human hepatocytes and human MASH liver specimens supported conservation of this signaling pathway.

**Conclusions:** These findings identify YY1 Ser247 phosphorylation as a critical molecular event linking non-apoptotic hepatocyte caspase-8 signaling to METRN induction and MASH fibrosis and define a previously unrecognized DED–SRC–GSK3β–YY1 signaling axis with potential therapeutic relevance.

## Introduction

Metabolic dysfunction-associated steatohepatitis (MASH) is a progressive form of metabolic dysfunction-associated steatotic liver disease (MASLD) and has become a leading cause of chronic liver disease worldwide[1, 2]. Progressive liver fibrosis is the histopathologic feature most strongly associated with liver-related and all-cause mortality in MASH, yet therapeutic options capable of preventing or reversing fibrosis remain limited[3–6]. The development of liver fibrosis is driven primarily by activation of collagen-producing hepatic stellate cells (HSCs)[3, 7], a process shaped by complex intercellular communication among hepatocytes, immune cells, endothelial cells and HSCs. Although hepatocytes have traditionally been viewed as the principal site of lipid accumulation and metabolic dysfunction during MASLD progression, increasing evidence indicates that they actively orchestrate fibrogenesis by secreting mediators that directly activate HSCs or remodel the hepatic microenvironment[8–11]. However, the hepatocyte-intrinsic signaling mechanisms that initiate these pro-fibrotic responses remain incompletely understood[11, 12].

Caspase-8 is best known as an initiator of death receptor-mediated apoptosis, but it can also perform non-apoptotic signaling functions in inflammation, differentiation, and tissue remodeling[13]. In MASH, hepatocyte apoptosis has long been considered a major pathogenic mechanism, prompting the development of pharmacological caspase inhibitors as potential therapies[14, 15]. However, drug-induced inhibition of caspase activity in a MASH clinical trial actually worsened liver fibrosis, suggesting that caspase-8 exerts biological functions beyond its canonical role in apoptosis[16]. Interestingly, genetic studies have similarly implicated caspase-8 in human liver disease, with variants at the *CASP8* locus associated with increased hepatic caspase-8 expression and chronically elevated plasma alanine aminotransferase concentrations[17, 18]. These combined observations suggest that understanding the non-apoptotic signaling functions of hepatocyte caspase-8 may provide new insight into the mechanisms driving liver fibrosis.

In this context, we recently identified a pathway in which hepatocyte caspase-8 promotes MASH fibrosis independently of apoptosis[19]. Using human liver specimens together with multiple diet-induced mouse models of MASH, we demonstrated that the non-apoptotic form of hepatocyte caspase-8 is markedly increased during MASH progression and that hepatocyte-specific deletion of caspase-8 suppresses hepatic stellate cell activation and liver fibrosis without altering hepatocyte apoptosis. Mechanistically, caspase-8 induced expression and secretion of the previously unrecognized HSC activator meteorin to promote liver fibrosis. Using cultured hepatocytes, we showed the involvement of the transcriptional regulator YY1 in *Metrn* induction, but the mechanism linking caspase-8 to YY1 and the role of YY1 in MASH fibrosis in vivo were not elucidated. Understanding how non-apoptotic caspase-8 can activate YY1 to induce pro-fibrotic meteorin is a critical goal, as it would add deep mechanistic insight and identify potentially new therapeutic targets in MASH fibrosis. Moreover, elucidating the caspase-8-YY1 link would contribute to our broader understanding of how caspase-8 can function in cells in a non-apoptotic manner and how cellular signaling pathways can regulate YY1, which is involved in many physiologic and pathophysiologic processes throughout the body[20].

Here we identify phosphorylation of YY1 at Ser247 as a critical regulatory event linking non-apoptotic hepatocyte caspase-8 signaling to MASH fibrosis. We show that the amino-terminal death effector domain (DED) of caspase-8 activates a Src–GSK3β signaling cascade that promotes YY1 Ser247 phosphorylation, nuclear accumulation, and transcriptional activation, leading to *Metrn* induction. Disruption of this pathway impairs YY1 activation, whereas phosphomimetic YY1 bypasses the requirement for hepatocyte caspase-8 and restores downstream fibrogenic signaling during MASH progression. Together, these findings identify the molecular switch controlling the caspase-8–YY1–meteorin fibrogenic pathway in MASH.

## Results

### Hepatocyte YY1 transduction in MASH mice increases *Metrn* expression and liver fibrosis in a caspase-8-dependent manner

We previously provided in vitro evidence that YY1 mediated the induction of *Metrn* downstream of caspase-8[19]. As a first step in testing a role for a caspase-8-YY1 axis in MASH fibrosis, we examined the effect of hepatocyte-specific YY1 transduction in wild-type and hepatocyte-caspase-8-knockout MASH mice. *Casp8^fl/fl^* mice were fed the fructose–palmitate–cholesterol (FPC) diet[9] for 10 weeks to establish steatosis and then injected the mice with AAV8-TBG-Cre to delete caspase-8 specifically in hepatocytes (Casp8-KO) or with control AAV8-TBG-GFP. In parallel, some of the control and Casp8-KO cohorts received AAV8-TBG-Yy1 to express YY1 in hepatocytes. The mice were then maintained on the FPC diet for an additional 6 weeks and then analyzed. The four experimental groups (control, YY1, Casp8-KO, Casp8-KO + YY1) exhibited comparable body weight, liver-to-body weight ratio, and blood glucose levels, indicating that neither hepatocyte-specific caspase-8 deletion nor YY1 overexpression altered systemic metabolic parameters (**Supplementary Fig. 1A-C**). We confirmed efficient deletion of hepatocyte caspase-8 and increased expression of YY1 in the Casp8-KO and YY1 groups, respectively (**Supplementary Fig. 1D-E**). Consistent with our previous study[19], hepatocyte-specific deletion of Casp8 reduced nuclear accumulation of YY1 and *Metrn* expression. Notably, YY1 overexpression increased both nuclear YY1 localization and *Metrn* expression in wild-type livers but failed to do so in hepatocyte-Casp8-knockout livers (**Fig.1A-B and Supplementary Fig. 1F**), indicating that YY1 activation is dependent on hepatocyte caspase-8 in vivo. Most importantly, AAV8-TBG-YY1 exacerbated fibrosis and HSC activation in control MASH mice, as evidenced by increased Sirius Red staining, elevated expression of fibrosis-associated genes (*Timp1, Spp1, Col1a1, Acta2*, and *Col1a2*), and enhanced α-SMA immunofluorescence, but failed to do so in hepatocyte-Casp8-KO mice (**Fig. 1C– E**). AAV8-TBG-YY1 also increased plasma ALT and hepatic mononuclear cells, a marker of myeloid-mediated liver inflammation, in control mice but not Casp8-KO mice (**Fig. 1F and Supplementary Fig. 1G,H**). Hepatic lipid accumulation was comparable among the four experimental groups (**Supplementary Fig. 1G,I**), indicating that these effects were not associated with changes in hepatosteatosis. Thus, hepatocyte-specific YY1 transduction increases *Metrn* expression, HSC activation, and liver fibrosis in experimental MASH in a caspase-8-dependent manner, indicating that caspase-8 is required to activate the pro-fibrotic YY1-meteorin pathway in vivo.

**Fig. 1.**
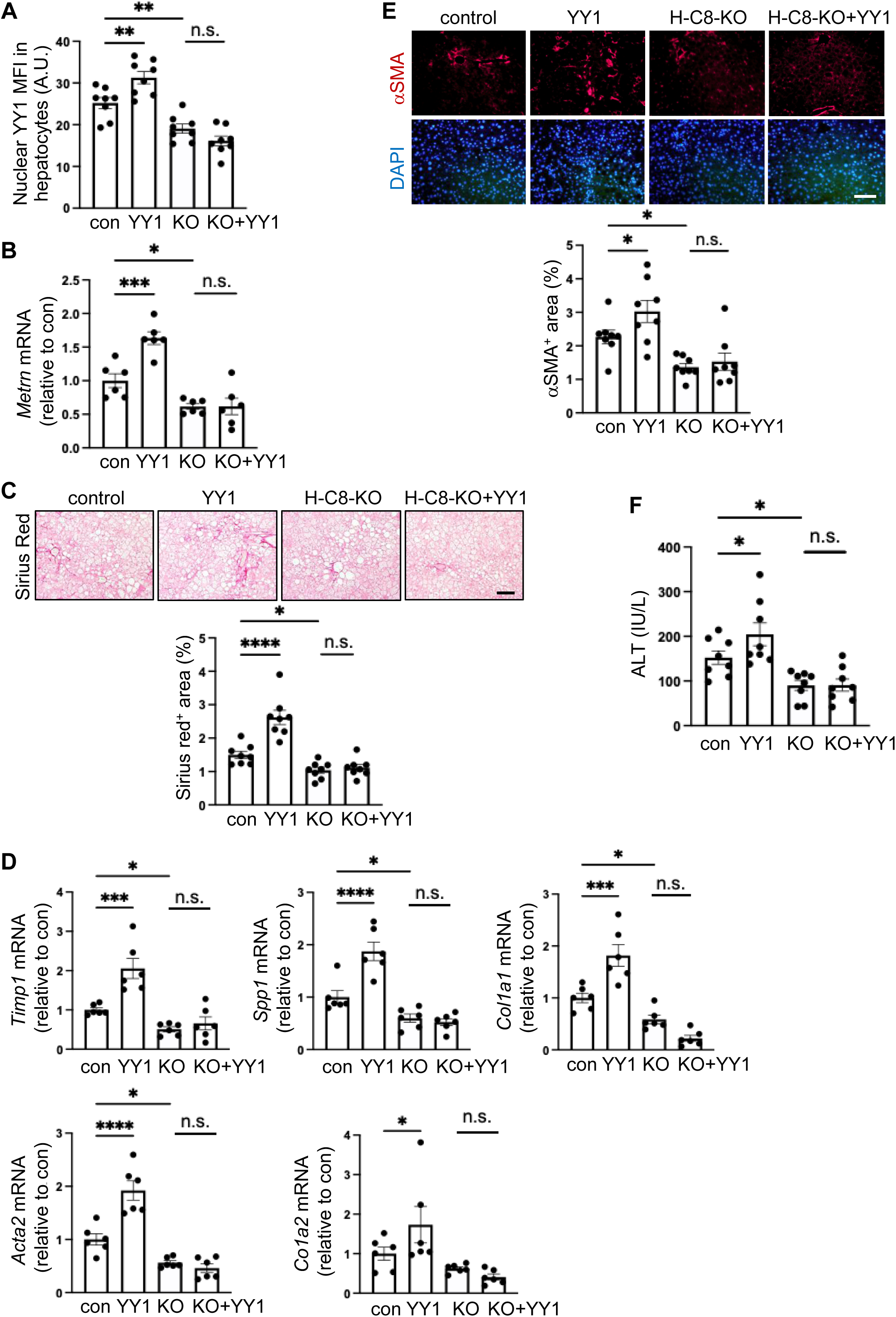
YY1 promotes liver fibrosis in wild-type mice but not hepatocyte-caspase-8-KO mice. *Casp8^fl/fl^* mice were fed the FPC diet for 10 wks, then injected with control AAV8-TBG-Cre (con), AAV8-TBG-Yy1 (YY1), AAV8-TBG-Cre to knockout hepatocyte caspase-8 (KO), or both the YY1 and Cre viruses (KO+YY1). The mice were then continued on the FPC diet for an additional 6 weeks. **A,** Nuclear YY1 MFI in hepatocytes as assessed by immunofluorescence microscopy (IFM). **B,** Hepatic *Metrn* mRNA. **C,** Staining of liver sections for Sirius red, with quantification of Sirius red-positive area; bar, 200 μm. **D,** Hepatic *Timp1*, *Spp1*, *Col1a1, Acta2*, and *Col1a2* mRNAs. **E,** α-SMA immunofluorescence (red) and quantification of α-SMA positive area, with nuclear DAPI counterstain (blue) shown in the bottom panels; bar, 200 μm. **F,** Plasma ALT. Data are presented as mean ± s.e.m.; *p < 0.05, **p < 0.01, ***p<0.001, ****p<0.0001, n.s., non-significant, one-way ANOVA; A, C, E, F, n = 8 per group. B, D, n = 6 per group.

### Caspase-8 promotes phosphorylation of YY1 at Ser247

The inability of hepatocyte YY1 transduction to restore *Metrn* expression or liver fibrosis in hepatocyte-Casp8-KO MASH mice suggested that hepatocyte caspase-8 regulates YY1 activity through a mechanism beyond controlling its expression. Because post-translational modification is a common mechanism regulating transcription factor activation, we next sought to identify YY1 post-translational modifications that might account for caspase-8-dependent YY1 activation. HA-tagged YY1 was expressed in Yy1-knockout AML12 hepatocytes, immunoprecipitated from nuclear extracts (**Supplementary Fig. 2A,B**), and analyzed by LC–MS/MS. Based on data analysis, we excluded oxidation, demethylation, glycosylation, and deamidation, but we discovered three potential YY1 phosphorylation sites (**Supplementary Fig. 2C**). Among the candidate phosphorylation sites identified, Ser247 exhibited the highest confidence based on localization probability, posterior error probability, and peptide intensity and was therefore selected for subsequent functional studies (**Supplementary Fig. 2D and Supplementary Table 1**).

To determine whether Ser247 represents a functionally important phosphorylation site, AML12 hepatocytes were transfected with HA-tagged wild-type YY1 or a phosphorylation-defective YY1-S247A mutant. Immunoprecipitation of HA-YY1 followed by phosphoserine immunoblotting demonstrated a marked reduction in YY1 serine phosphorylation in cells expressing YY1-S247A compared with wild-type YY1, indicating that Ser247 is a major phosphorylation site in hepatocytes (**Fig. 2A**). Likewise, partial silencing of Casp8 substantially reduced YY1 serine phosphorylation (**Fig. 2B**), demonstrating that phosphorylation of YY1 depends on hepatocyte caspase-8. Consistent with the reduction in YY1 phosphorylation following Casp8 silencing, HA-tagged YY1 showed lower nuclear localization in Casp8-knockout vs. wild-type AML12 hepatocytes (**Fig. 2C**), suggesting that caspase-8-dependent phosphorylation promotes YY1 nuclear localization. We next examined whether Ser247 phosphorylation influences YY1 subcellular localization and transcriptional activity. Whereas wild-type YY1 accumulated predominantly in the nucleus, the phosphorylation-defective YY1-S247A mutant exhibited markedly reduced nuclear localization (**Fig. 2D**). Consistent with this observation, expression of wild-type YY1 significantly increased *Metrn* mRNA, whereas YY1-S247A failed to induce Metrn above basal levels (**Fig. 2E**), indicating that Ser247 phosphorylation is required for YY1-mediated induction of *Metrn*. To further determine whether Ser247 phosphorylation is sufficient to activate YY1 independently of caspase-8, we transfected Casp8-KO AML12 hepatocytes with a phosphomimetic mutant in which Ser247 was replaced with aspartate (YY1-S247D). YY1-S247D cause an approximately two-fold greater nuclear accumulation than wild-type YY1 in the Casp8-KO hepatocytes (**Fig. 2F**), and YY1-S247D, but not wild-type YY1, increased *Metrn* expression above the basal level in these cells (**Fig. 2G**). Finally, to determine whether the caspase-8-dependent regulation of YY1 phosphorylation observed in cultured hepatocytes also occurs in vivo, YY1 was immunoprecipitated from liver lysates of FPC-fed MASH mice and analyzed by phosphoserine immunoblotting. Phosphorylated YY1 was readily detected in wild-type MASH livers and showed an increase following hepatocyte YY1 transduction. In contrast, hepatocyte-specific deletion of Casp8 reduced YY1 serine phosphorylation, and YY1 overexpression failed to restore phosphorylation in the hepatocyte-Casp8-KO mice **(Fig. 2H**). These findings demonstrate that phosphorylation of YY1 at Ser247 is necessary for YY1 activation in wild-type hepatocytes and is able to restore YY1-dependent *Metrn* induction in the absence of hepatocyte caspase-8, establishing Ser247 phosphorylation as a critical molecular switch linking caspase-8 to YY1 activation.

**Fig. 2.**
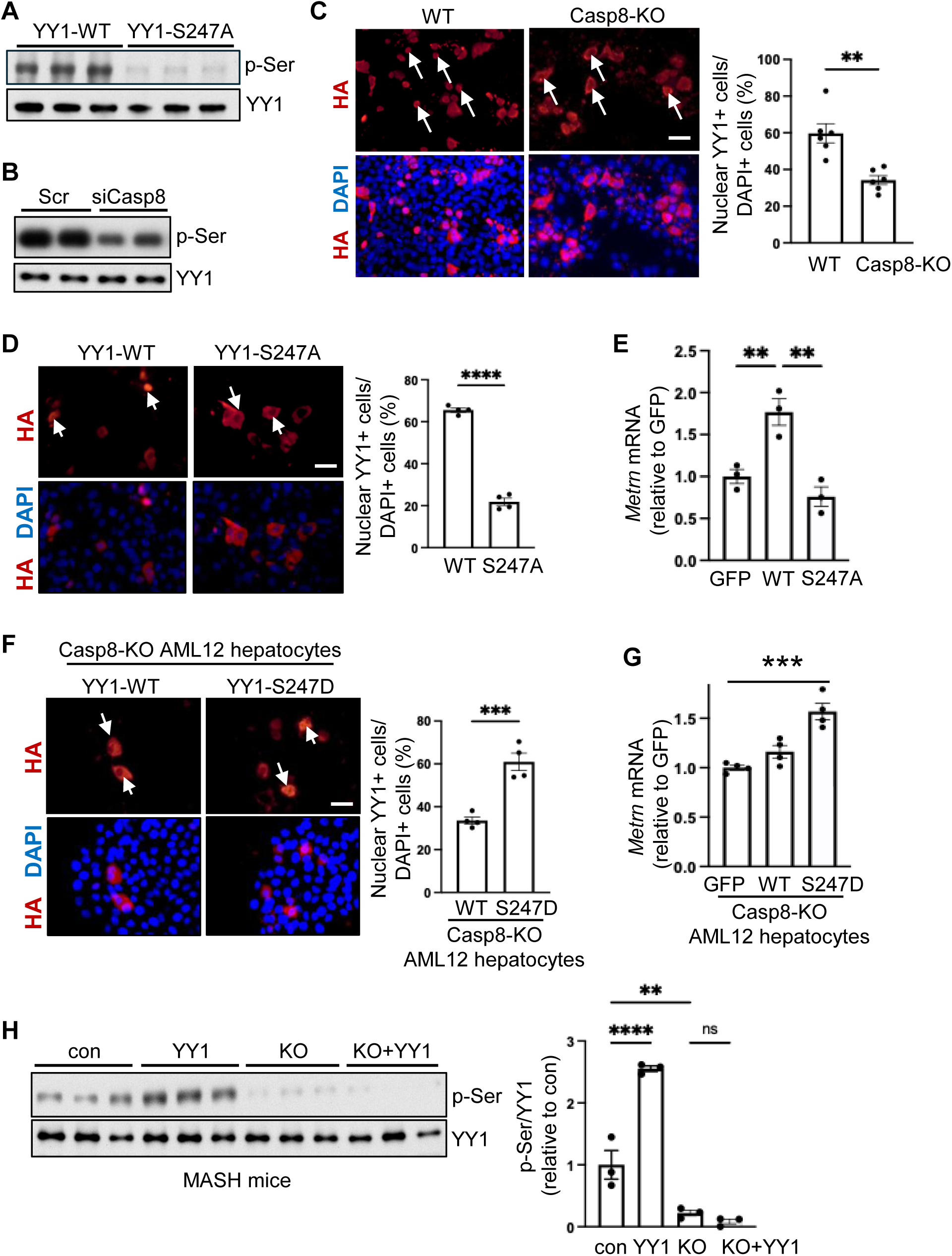
Caspase-8 promotes Ser247-phosphorylation of YY1. **A,** AML12 hepatocytes were transfected with wild-type HA-tagged YY1 (YY1-WT) or YY1-S247A. HA-YY1 was immunoprecipitated (IP) and immunoblotted for phosphoserine (p-Ser) and YY1. **B,** AML12 hepatocytes were treated with control scrambled RNA (Scr) or siCasp8 and then analyzed as in panel A. **C,** Wild-type (WT) or caspase-8-KO AML12 hepatocytes expressing HA-YY1 were quantified by IFM for nuclear HA-YY1-positive cells. Unpaired two-tailed Student’s t-test; n = 6/group. Scale bar, 100 μm. **D,** AML12 hepatocytes expressing HA-YY1-WT or HA-YY1-S247A were analyzed as in panel c. Unpaired two-tailed Student’s t-test; n = 4 per group. Scale bar, 20 μm. **E,** *Metrn* mRNA expression was measured in AML12 hepatocytes expressing GFP, YY1-WT, or YY1-S247A. One-way ANOVA; n = 3 per group. **F,** Caspase-8-KO AML12 hepatocytes expressing HA-YY1-WT or HA-YY1-S247D were analyzed by IFM for nuclear HA-YY-positive cells. Unpaired two-tailed Student’s t-test; n = 4 per group. Scale bar, 20 μm. **G,** *Metrn* mRNA was measured in caspase-8-KO AML12 hepatocytes expressing GFP, HA-YY1-WT, or HA-YY1-S247D. One-way ANOVA; n = 4 per group. **H,** YY1 was immunoprecipitated from liver lysates of the mice shown in Fig. 1 and immunoblotted for p-Ser and YY1. Densitometric quantification of the p-Ser/YY1 ratio is shown. Data are presented as mean ± s.e.m.; **p<0.01, ***p<0.001, ****p<0.0001.

### Constitutively active YY1 promotes MASH fibrosis in the absence of hepatocyte caspase-8

To translate the above in vitro findings to MASH, we tested the prediction that the transduction of hepatocytes with constitutively active S247D YY1 would exacerbate liver fibrosis in MASH mice lacking hepatocyte caspase 8. Accordingly, *Casp8^fl/fl^* mice were fed the FPC diet for 10 weeks and then injected with AAV8-TBG-Cre to delete hepatocyte caspase-8. At the same time, the mice were administered AAV8 vectors expressing either wild-type YY1 or YY1-S247D mutant. The mice were analyzed after 6 additional weeks on the FPC diet. The wild-type and YY1-S247D transgene caused a similar increase in hepatic *Yy1* mRNA expression (**Supplementary Fig. 3A**). Body weight, liver-to-body weight ratio, and blood glucose were comparable among the four experimental groups (**Supplementary Fig. 3B-D**). Consistent with our in vitro findings, expression of YY1-S247D, but not wild-type YY1, restored nuclear accumulation of YY1 in hepatocyte-caspase-8-deficient livers (**Fig. 6A, Supplementary Fig. 3E**). Likewise, YY1-S247D restored hepatic *Metrn* expression to levels approaching those observed in wild-type mice, whereas wild-type YY1 remained largely ineffective (**Fig. 3B**). Most importantly, YY1-S247D, but not wild-type YY1, restored liver fibrosis in the hepatocyte-caspase-8-KO MASH mice as demonstrated by increased Sirius Red staining (**Fig. 3C**) and elevated expression of fibrosis-associated genes, including *Timp1, Col1a1, Tgfb1,* and *Acta2* (**Fig. 3D**). Similar findings were observed using the endpoints of α-SMA-positive area, which is a marker of activated HSCs, and hepatic COL1A1 deposition (**Fig. 3E,F**). These changes occurred despite similar hepatic lipid accumulation among the four experimental groups (**Supplementary Fig. 3F,G**). We also found that YY1-S247D restored hepatic mononuclear cell infiltration, F4/80-positive macrophage accumulation, *Mcp1* expression, and plasma ALT in hepatocyte-Casp8-deficient mice (**Fig. 3G and Supplementary Fig. 3H–J**), whereas wild-type YY1 had little effect on any of these parameters. These data support the importance of casapase-8-dependent phosphorylation of YY1 at S247 in MASH progression.

**Fig. 3.**
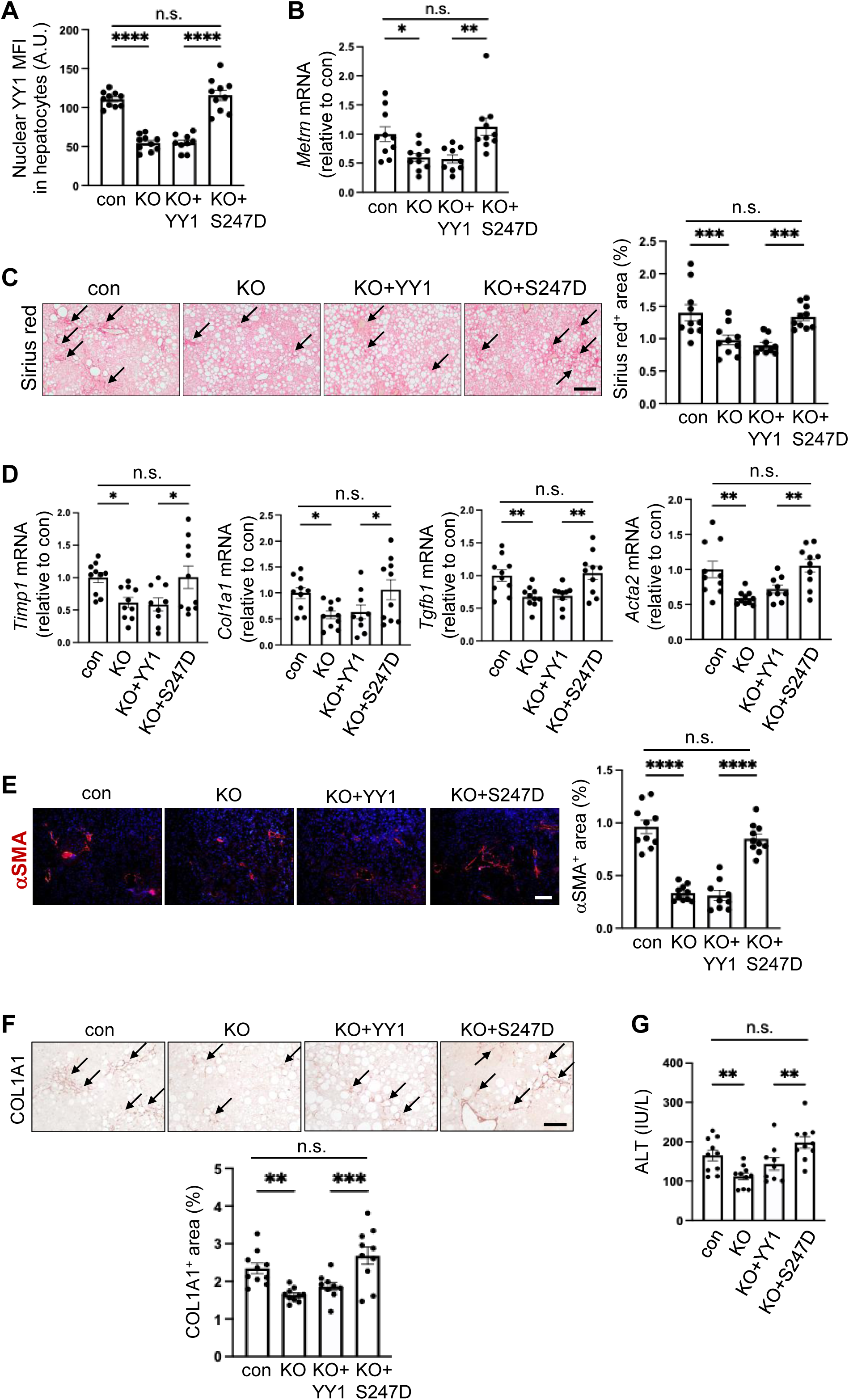
Constitutively active S247D-YY1 exacerbates liver fibrosis in hepatocyte-specific caspase-8-deficient MASH mice. *Casp8^fl/fl^* mice were fed the FPC diet for 10 wks, then injected with control AAV8-TBG-GFP (con), AAV8-TBG-Cre to knockout hepatocyte caspase-8 (KO), both AAV8-TBG-Cre and AAV8-TBG-YY1 (KO+YY1), or AAV8-TBG-Cre and AAV8-TBG-S247D-YY1 (KO+S247D). The mice were then continued on the FPC diet for an additional 6 weeks. **A,** nuclear YY1 MFI in hepatocytes as assessed by IFM. **B,** Hepatic *Metrn* mRNA. **C,** Staining of liver sections for Sirius red, with quantification of Sirius red-positive area (arrows in images); bar, 500 μm. **D,** Hepatic *Timp1, Col1a1*, *Tgfb1*, and *Acta2* mRNAs. **E,** α-SMA immunofluorescence (red) and quantification of α-SMA positive area; bar, 200 μm. **F,** COL1A1 immunohistochemistry, with quantification of COL1A1 positive area; bar, 200 μm. **G,** Plasma ALT. a-g, Data are presented as mean ± s.e.m.; one-way ANOVA; con, n = 10, KO, n = 10, KO+YY1, n = 9, KO+S247D, n = 10. *p < 0.05, **p < 0.01, ***p<0.001, ****p<0.0001, n.s., non-significant.

### Src links caspase-8 to YY1 activation in hepatocytes

Identification of Ser247 as the critical phosphorylation site on YY1 prompted us to investigate the signaling pathway linking hepatocyte caspase-8 to YY1 activation. Because caspase-8 lacks intrinsic kinase activity, we reasoned that an intermediary kinase must mediate YY1 phosphorylation. Previous studies have implicated non-apoptotic caspase-8 signaling in Src activation in several cell types[21]. (Although Src is a tyrosine kinase, we reasoned that if Src were involved, it could secondarily activate a serine kinase to link to phospho-Ser-YY1, as described below.) Our initial studies showed that partial silencing of caspase-8 reduced phosphorylation of Src at its activating Y416 residue while having little effect on total Src abundance (**Fig. 4A**), consistent with caspase-8-mediated Src activation. Most importantly, partial Src silencing decreased both nuclear YY1 (**Fig. 4B**) and *Metrn* expression (**Fig. 4C**), demonstrating that Src activity contributes to YY1-dependent induction of *Metrn*. To determine whether engineered Src activation was sufficient to overcome the absence of hepatocyte caspase-8, control and caspase-8-silenced AML12 hepatocytes were transduced with either wild-type Src or a constitutively active Src mutant (Y418D). The data show that Y418D-Src, but not wild-type Src, restored both nuclear YY1 activity and *Metrn* expression in these caspase-8-silenced hepatocytes (**Fig. 4D,E**). Thus, caspase-8-mediated Src plays a key role in YY1-mediated *Metrn* induction in hepatocytes.

**Fig. 4.**
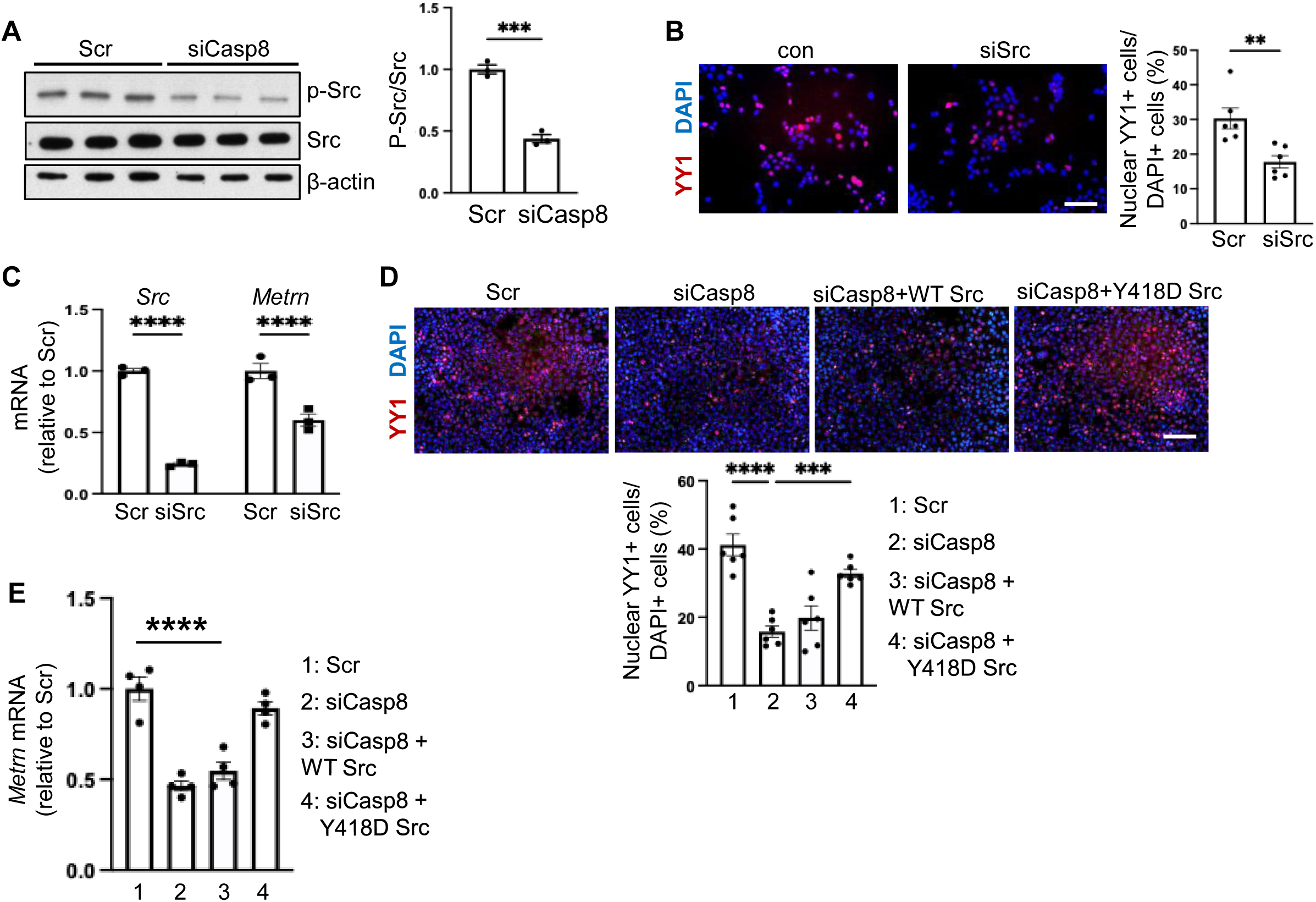
Src links caspase-8 to the activation of YY1-mediated induction of meteorin. **A,** Immunoblot and quantification of p-Y416-Src and total Src in control (Scr) and caspase-8-silenced AML12 hepatocytes. **B,** Nuclear YY1-positive cells as assessed by IFM in control (Scr) and Src-silenced AML12 hepatocytes. Scale bar, 100 μm. *n* = 4 biologically independent samples per group. Statistical significance was determined by two-tailed unpaired Student’s *t*-test. **C,** *Src* and *Metrn* mRNA in control (Scr) and Src-silenced AML12 hepatocytes. n = 3 biologically independent samples per group. Statistical significance was determined by two-tailed unpaired Student’s *t*-test. **D,** Nuclear YY1-positive cells as assessed by IFM in control (Scr) and caspase-8-silenced AML12 hepatocytes transduced with wild-type Src or constitutively activeY418D-Src. Scale bar, 200 μm. *N* = 6 biologically independent samples per group. Statistical significance was determined by one-way ANOVA. **E,** *Metrn* mRNA in control (Scr) and caspase-8-silenced AML12 hepatocytes transduced with wild-type Src or Y418D-Src. n = 4 biologically independent samples per group. Statistical significance was determined by one-way ANOVA. Data are presented as mean ± s.e.m. **p<0.01, ***p<0.001, ****p<0.0001.

### GSK3β links Src activation to YY1 phosphorylation

As noted above, we reasoned that an intermediary serine/threonine kinase must couple Src tyrosine kinase activation to YY1 serine phosphorylation. To identify this kinase, we made a list of serine kinases predicted to regulate YY1 activity and that were possible or known targets of Src and then performed a candidate siRNA screen for *Metrn* expression in AML12 hepatocytes. Among the candidates examined, GSK3β and ACVR2A produced the greatest reduction in *Metrn* expression (**Supplementary Fig. 4A**). However, whereas silencing ACVR2A did not alter nuclear localization of YY1 (**Supplementary Fig. 4B**), GSK3β knockdown reduced both the nuclear accumulation of YY1 (**Fig. 5A**) and *Metrn* expression (**Fig. 5B**). Moreover, silencing either caspase-8 or Src reduced Y216-phosphorylation/activation of GSK3β (**Fig. 5C**), indicating that GSK3β is activated by both of these upstream proteins. Most importantly, silencing either Src or GSK3β in AML12 hepatocytes suppressed *Metrn* expression (**Fig. 5D**). These combined data support a caspase-8-Src-GSK3β-YY1-meteorin pathway in hepatocytes. To determine whether this pathway is conserved in primary hepatocytes, we repeated these studies in freshly isolated mouse hepatocytes. Similar to AML12 hepatocytes, Src silencing reduced phosphorylation of GSK3β (**Fig. 5E and Supplementary Fig. 4C**), and knockdown of either Src or GSK3β significantly decreased *Metrn* expression (**Supplementary Fig. 4D**). We next extended these observations to primary human hepatocytes. Consistent with the mouse studies, caspase-8 silencing reduced phosphorylation of Src and GSK3β (**Fig. 5f**) and *METRN* expression (**Fig. 5G**), demonstrating conservation of the Src–GSK3β–YY1 signaling pathway across species. Finally, we investigated whether this signaling pathway is altered in human MASH. Immunoblot analysis of human liver specimens demonstrated increased Src phosphorylation in MASH (**Fig. 5H**), and primary hepatocytes isolated from MASH livers exhibited corresponding increases in the p-Src/Src and p-GSK3β/GSK3β ratios (**Fig. 5I**), supporting the relevance of the signaling pathway to human MASH.

**Fig. 5.**
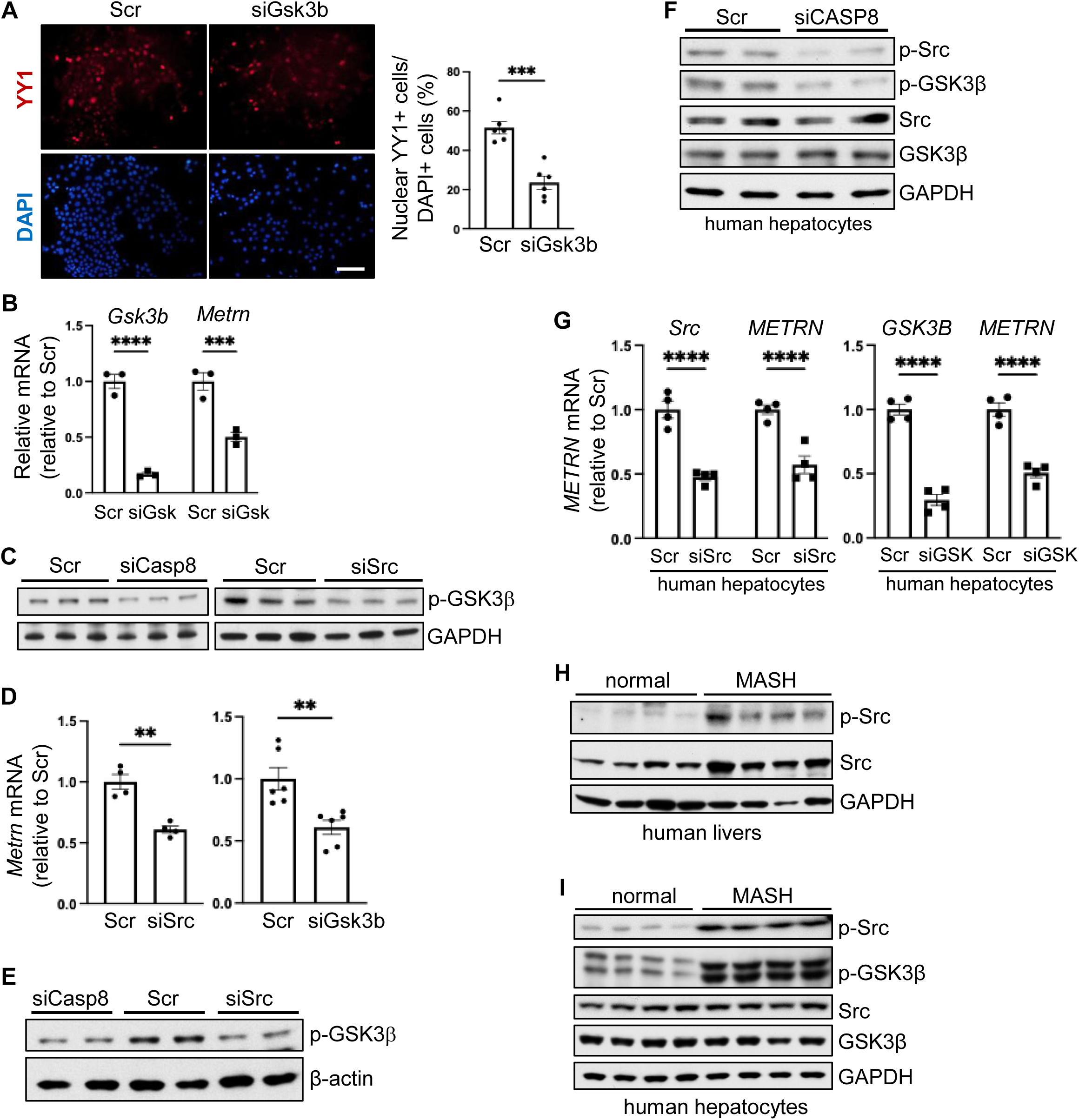
GSK3β regulates nuclear YY1 downstream of Src. **A,** Nuclear YY1-positive cells as assessed by IFM in control (Scr) and GSK3β-silenced AML12 hepatocytes. Scale bar, 100 μm. n = 6 biologically independent samples per group. Statistical significance was determined by two-tailed unpaired Student’s *t*-test. **B,** *Gsk3b* and *Metrn* mRNA in control (Scr) and GSK3β-silenced (siGsk) AML12 hepatocytes. n = 3 biologically independent samples per group. Statistical significance was determined by two-tailed unpaired Student’s t-test. **C,** Immunoblot of p-Y216-GSK3β in control (Scr), Src-silenced, and Casp8-silenced AML12 hepatocytes. **D,** *Metrn* mRNA in control, Src-silenced, and GSK3β-silenced primary mouse hepatocytes. siSrc data, n = 4; siGsk3b data, n = 6 biologically independent samples per group. Statistical significance was determined by two-tailed unpaired Student’s *t*-test. **E,** Immunoblot of p-Y216-GSK3β in control and Src-silenced primary mouse hepatocytes. **F,** Immunoblot of phospho- and total Src and GSK3β in control (Scr) and caspase-8-silenced primary human hepatocytes. **G,** *METRN* mRNA in control (Scr)-treated and Src- and GSK3β-silenced primary human hepatocytes. n = 4 biologically independent samples per group. Statistical significance was determined by two-tailed unpaired Student’s t-test. **H,** Immunoblot of p-Src and total Src in normal and MASH human liver samples. **i,** Immunoblot of phospho- and total Src and GSK3β in normal and MASH human hepatocytes. Data are presented as mean ± s.e.m. **p<0.01, ***p<0.001, ****p<0.0001.

### The DED domain of caspase-8 activates the Src–GSK3β–YY1 signaling pathway

Having established that Src and GSK3β mediate YY1 activation downstream of hepatocyte caspase-8, we next sought to determine how caspase-8 initiates this kinase cascade. Previous studies have shown that the amino-terminal death effector domains (DEDs) of caspase-8 mediate non-apoptotic signaling through protein–protein interactions independently of the catalytic domain[22], and in our previous study, we showed that the caspase-8 DED domain could restore *Metrn* expression in caspase-8-silenced hepatoctyes in vitro[19]. Moreover, the caspase-8 DED domain has been shown to interact with Src in cancer cells and 293T cells[22, 23]. We therefore investigated whether the DED domain functions as the signaling module responsible for activation of the Src–GSK3β–YY1-metoerin pathway in hepatocytes. We first examined whether endogenous caspase-8 associates with Src in hepatocytes. Immunoprecipitation of Src from wild-type and caspase-8-KO AML12 hepatocytes revealed co-immunoprecipitation of caspase-8 with Src in wild-type hepatocytes, and phosphorylated Src was readily detected in the Src immunoprecipitates from wild-type hepatocytes but was markedly reduced in caspase-8-KO hepatocytes (**Fig. 6A**). These data are consistent with the idea hepatocyte caspase-8 promotes Src activation through caspase-8-Src association. Previous structural studies identified Asp73 within the conserved RXDLL motif of the caspase-8 DED domain as a critical determinant of Src binding[22, 23]. To determine whether this motif is required for activation of the downstream signaling pathway, wild-type caspase-8 or the interaction-defective caspase-8-D73A mutant was expressed in caspase-8-KO AML12 hepatocytes. Whereas wild-type caspase-8 restored phosphorylation of both Src and GSK3β in the KO hepatocytes, the D73A mutant failed to rescue activation of either kinase (**Fig. 6B**). Consistent with these findings, genetically induced expression of wild-type caspase-8 in the KO cells restored nuclear accumulation of YY1 (**Fig. 6C**) and *Metrn* expression (Fig. **6D**), whereas the caspase-8-D73A mutant exhibited little activity. These data demonstrate that the conserved RXDLL motif is essential for activation of the Src–GSK3β–YY1-*Metrn* signaling pathway downstream of caspase-8.

**Fig. 6.**
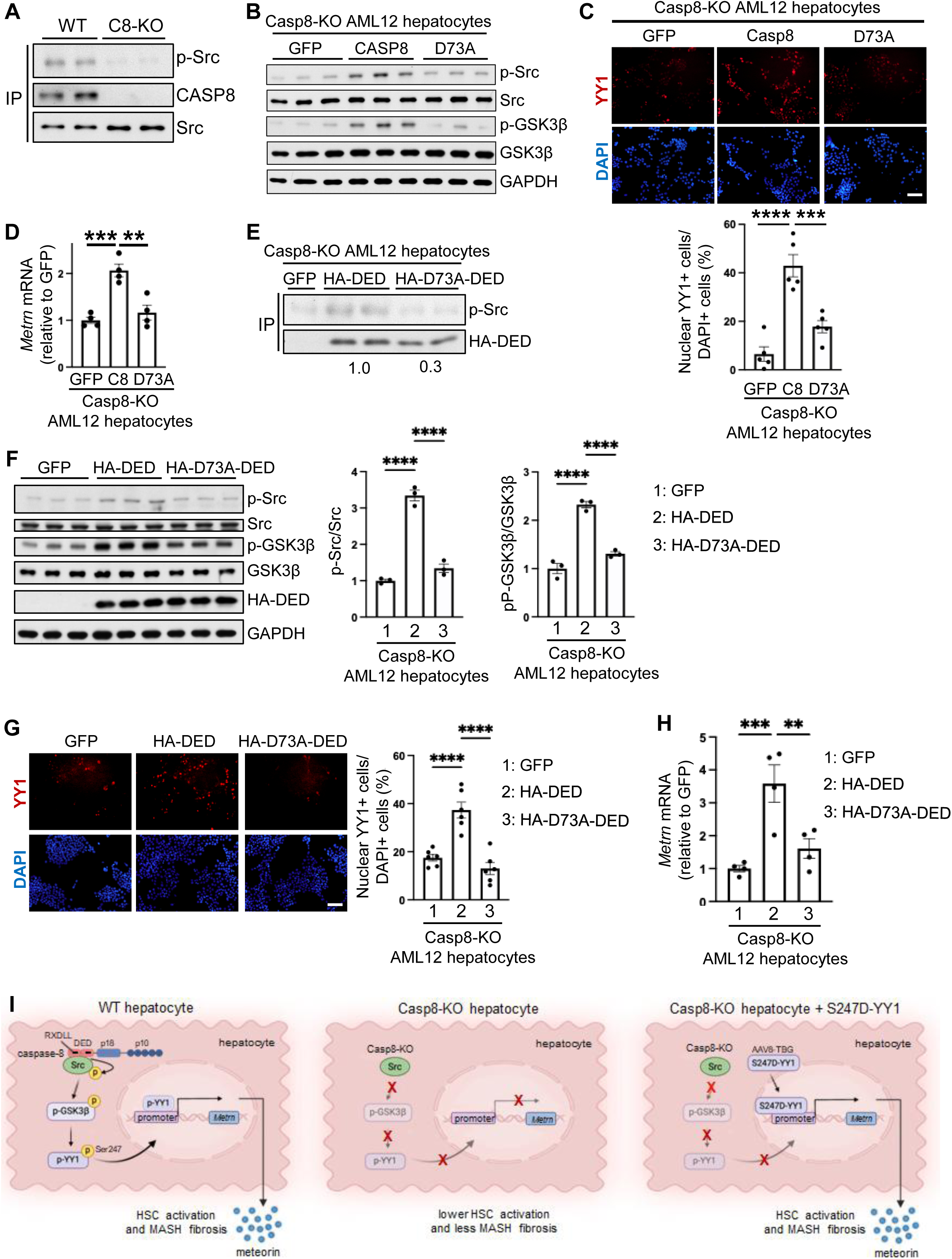
The DED domain of caspase-8 regulates Src activation and nuclear YY1 localization. **A,** Src was immunoprecipitated (IP) from wild-type and caspase-8-KO AML12 hepatocytes and immunoblotted for p-Y416-Src, caspase-8, and total Src. **B,** Immunoblot analysis of phosphorylated and total Src and GSK3β in wild-type and caspase-8-KO AML12 hepatocytes transfected with control GFP, wild-type caspase-8, or D73A-caspase-8. **C,** IFM images and quantification of nuclear YY1-positive cells in caspase-8-KO AML12 hepatocytes transfected as in panel B. Scale bar, 200 μm. n = 5 biologically independent experiments per group. Statistical significance was determined by one-way ANOVA. **D,** *Metrn* mRNA in caspase-8-KO AML12 hepatocytes transfected as in panel B. *n* = 4 biologically independent experiments per group. Statistical significance was determined by one-way ANOVA. **E,** HA-tagged proteins were immunoprecipitated from caspase-8-KO AML12 hepatocytes transfected with GFP, HA-DED, or HA-D73A-DED and then immunoblotted for p-Src and HA. Densitometric quantification of the p-Ser/HA-DED is shown below the blot. **F,** Immunoblot and quantification of phospho-and total Src and GSK3β and HA-DED in caspase-8-KO AML12 hepatocytes transfected as in panel E. The data are normalized for the GFP-transfected control cells. n = 3 biologically independent experiments per group. Statistical significance was determined by one-way ANOVA. **G,** IFM images and quantification of nuclear YY1-positive cells in caspase-8-KO AML12 hepatocytes transfected as in panel E. Scale bar, 200 μm. n = 6 biologically independent experiments per group. Statistical significance was determined by one-way ANOVA. **H,** *Metrn* mRNA in caspase-8-KO AML12 hepatocytes transfected as in panel E. n = 4 biologically independent experiments per group. Statistical significance was determined by one-way ANOVA. Data in panels A-H are presented as mean ± s.e.m. **p<0.01, ***p<0.001, ****p<0.0001. **I**, Proposed model of the hepatocyte caspase-8–SRC–GSK3β–YY1-*Metrn* signaling pathway in MASH, integrated with our previous study showing that caspase-8 is upregulated in human and experimental MASH, which leads to increased meteorin and meteorin-induced HSC activation and liver fibrosis[19]. *Left*, In MASH hepatocytes, the DED domain of caspase-8, through its RXDLL motif, promotes SRC phosphorylation/activation, which then phosphorylates/activates GSK3β. Activated GSK3β leads to YY1 phosphorylation at Ser247, causing YY1 nuclear accumulation, YY1-mediated induction of *Metrn*, and meteorin-mediated HSC activation/liver fibrosis. *Middle*, When hepatocyte caspase-8 is deleted in MASH, the Src-GSK3β-YY1-*Metrn* pathway becomes inactive, resulting in lower meteorin production and a decrease in HSC activation/liver fibrosis. *Right*, meteorin expression and liver fibrosis can be restored in hepatocyte-caspase-8-KO MASH mice through in vivo transduction of hepatocytes with phosphomimetic S247D-YY1 (AAV8-TBG-S247D-YY1). The image was created with BioRender.com (agreement # MU2A5RNGGB; https://BioRender.com/0gclsza).

We next asked whether the DED domain alone is sufficient to activate this pathway. caspase-8-KO AML12 hepatocytes were transfected with HA-tagged wild-type DED or D73A-DED constructs. Immunoprecipitation of HA-tagged proteins demonstrated robust association of the wild-type DED with activated Src, whereas the D73A mutation markedly reduced this interaction (**Fig. 6E**). Further, expression of wild-type DED, but not D73A DED, was sufficient to restore phosphorylation of Src and GSK3β in Casp8-deficient cells (**Fig. 6F**), nuclear YY1 (**Fig. 6G and Supplementary Fig. 5A**), and Metrn expression (**Fig. 6H**). Note that the DED domain does not interact directly with YY1 (**Supplementary Fig. 5B**). These combined data, when considered with our previous findings[19], support a mechanism in which the DED-RXDLL motif of caspase-8 in hepatocytes activates a Src–GSK3β kinase pathway, leading to phosphorylation-induced activation of YY1 and then YY1-mediated induction of pro-fibrotic meteorin (**Fig. 6I**).

## Discussion

The findings in this report define a previously unreported caspase-8-DED–Src–GSK3β–phospho-YY1 signaling pathway in hepatocytes that, via meteorin induction, is relevant to the most clinically important feature of MASH, liver fibrosis. This work addresses the key mechanistic question arising from our recent caspase-8-MASH study[19], namely, identifying how a non-apoptotic function of caspase-8 can promote meteorin-induced HSC activation and liver fibrosis in MASH. Moreover, our discovery that phosphorylation of YY1 is a key step in the pathway contributes new insights into how this universally important transcriptional regulator can be regulated. Previous in vitro studies have shown that YY1 can be phosphorylated at multiple sites[24–31]. One of these studies presented evidence that endothelial pS118-YY1 plays a role in atherosclerosis[31], while the others focused on correlations between YY1 phosphorylation and the stage of the cell cycle and/or on the role of YY1 phosphorylation in either increasing or decreasing YY1-DNA interaction in vitro. Interestingly, one study showed that T-A mutations at T348 and T378 in the DNA-binding domain of YY1 abolished YY1-DNA interaction, whereas neither S247A nor S247D in the spacer domain of YY1 affected this endpoint[24]. Here, using in vivo transduction of hepatocytes with wild-type YY1 or phosphomimetic S247D-YY1 in MASH mice lacking hepatocyte-caspase-8, we show that S247-YY1 phosphorylation links caspase-8 to the induction of *Metrn* and liver fibrosis in MASH.

Apart from the role of YY1 phosphorylation per se, a previous study showed that overall hepatic YY1 expression as assessed by immunoblot in whole liver extracts was increased in mice and humans with metabolic liver disease-associated hepatosteatosis[32]. The investigators also showed that overexpressing YY1 using adeno-Yy1 promoted steatosis, whereas adeno-shYy1 reduced hepatic triglyceride accumulation. That authors linked this effect to YY1-mediated repression of farnesoid X receptor (FXR) through YY1 binding within intron 1 of the *FXR* locus[32]. Our study, by administering AAV8-TBG-Yy1 or -S247D-Yy1 after diet-induced steatosis had already developed, focused on the progression of hepatosteatosis to fibrotic MASH, which is the clinically most important stage in metabolic liver disease. Although it is theoretically possible that caspase-8-mediated phosphorylation/activation of YY1 might activate FXR, none of our caspase-8 or YY1 interventions reversed hepatosteatosis.

Our previous study showed that the role of caspase-8 in the meteorin-MASH fibrosis pathway was independent of its apoptotic protease activity[19], and the current data shows how this occurs, i.e., by DED-mediated Src activation. Our work in this area was inspired by previous studies investigating the role of the caspase-8-DED domain in cancer cell migration[23]. In invasive tumor cells, deletion of the DED-RXDLL impairs Src-mediated signaling and migration[23]. These data suggest that DED-RXDLL motif in caspase-8 can regulate the assembly of caspase-8-containing signaling complexes, including those involving Src. This idea is consistent with our findings that the wild-type DED, but not D73A-DED, associated with p-Src and was sufficient to activate Src– GSK3β–YY1 signaling in hepatocytes.

Our finding of a non-apoptotic role of caspase-8 in MASH progression is supported by several pieces of human data, including data from analyses of human MASH liver[19] and from studies with human hepatocytes[19], human genetic data[17, 18], and clinical trial data showing that drug-induced inhibition of pro-apoptotic caspase activity did not improve liver fibrosis in MASH subjects[16]. However, the pro-apoptotic function of caspase-8 could be relevant to other types of liver disease. In this context, a recent study reported that hepatocyte-specific loss of JNK1 and JNK2 markedly increased oxidative stress, apoptosis, liver injury and fibrosis, and that inhibition of caspase-8 in this model ameliorated these phenotypes[33]. The authors concluded that, in the setting of hepatocyte JNK1/2 deficiency, caspase-8 primarily mediates apoptosis-associated liver injury. To place these findings in context, JNK1/2-deficient mice represent a genetically engineered, highly apoptosis-prone model in which hepatocyte JNK signaling is deleted early in life. Moreover, caspase-8 inhibition in this model reduced hepatosteatosis, suggesting complicated effects on liver metabolism. Nonetheless, this interesting study suggests that caspase-8-mediated apoptosis could play a role in diseases in which hepatocyte apoptosis plays a major role in liver injury.

The mechanistic discoveries in this study have potential therapeutic implications for MASH progression. By providing deeper mechanistic support for the meteorin pathway, the findings here strengthen the premise for developing therapies that genetically silence hepatocyte meteorin, including through the use of GalNAc-conjugated oligonucleotides[34]. However, it is possible that YY1 promotes MASH progression by additional mechanisms, and thus targeting the DED–Src– GSK3β–YY1 signaling axis may be more potent in preventing MASH progression than silencing only meteorin. Although direct pharmacological targeting of an individual YY1 phospho-site is challenging, emerging approaches that exploit site-specific docking interactions or targeted dephosphorylation[35, 36] may ultimately provide strategies to block YY1-Ser247-phosphorylation to block YY1-mediated MASH progression. It may also be possible to selectively target upstream components in the pathway in hepatocytes while limiting systemic effects by using hepatocyte-directed therapeutic platforms.

Our study has several limitations. Although our data position GSK3β upstream of YY1-Ser247 phosphorylation, they do not establish whether GSK3β directly phosphorylates this residue or regulates its phosphorylation indirectly through additional signaling molecules. Moreover, as alluded to above, future work will need to explore whether additional YY1 target genes besides *METRN* contribute to MASH progression. The structural consequences of Ser247 phosphorylation and the mechanisms by which this modification promotes YY1 nuclear accumulation also remain to be defined. Finally, the metabolic or inflammatory signals responsible for the increase in hepatocyte caspase-8 during MASH remain unknown. Addressing these questions will further refine our understanding of the hepatocyte signaling networks that drive MASH progression, notably liver fibrosis, and may reveal new opportunities for therapeutic intervention.

## Methods

This research complies with all relevant ethical regulations and has been approved by Columbia University’s Institutional Animal Care and Use Committee (IACUC) and Institutional Review Board.

### Experimental animals

Animal protocols used were approved by Columbia University’s Institutional Animal Care and Use Committee (protocol no. AC-AABL5573) and were cared for according to institutional and National Institutes of Health (NIH) guidelines for the care and use of laboratory animals. The mice were socially housed in standard cages at 22^°^C and with a 12-hour light-dark cycle in a barrier facility with ad libitum access to food and water. Male wild-type C57BL/6J mice (#000664) and *Casp8^fl/fl^*mice (#027002, on C57BL/6J background[37]) were obtained from Jackson Laboratory (Bar Harbor, ME) and acclimated for one week at the Columbia University Irving Medical Center Institute of Comparative Medicine before being randomly assigned to experimental groups. To induce MASH, the mice were fed a fructose-palmitate diet with 1.25% cholesterol (FPC diet, Enotiv, TD.160785 PWD) for 16 weeks to induce MASH[9, 38]. The chow diet was obtained from LabDiet (catalog #5053). For hepatocyte-specific gene manipulation, Casp8*^fl/fl^* mice were maintained on the FPC diet for 10 weeks before receiving tail-vein injections of AAV8-TBG-Cre, AAV8-TBG-GFP, AAV8-TBG-YY1, or AAV8-TBG-YY1-S247D, as indicated in the corresponding figure legends. Each virus was administered at 2 × 10¹¹ genome copies per mouse, and the mice were kept on the FPC diet for an additional 6 weeks before analysis.

### Human liver specimens and human primary liver cells

De-identified human liver specimens were obtained from the Liver Tissue Cell Distribution System at the University of Minnesota. Both male and female specimens were collected at the time of liver transplantation and preserved as frozen samples. The written informed consent was obtained from all donors for tissue samples. Diagnostic information is provided in Supplementary Table 1. Phenotypic and pathological assessments were conducted by medical physicians and pathologists affiliated with the Liver Tissue Cell Distribution System. The study protocol was approved by the Institutional Review Board (IRB) at Columbia University Irving Medical Center. Diseased (MASH, METAVIR Score = F4) adult human liver cells were obtained from the Organ Processing Hub and the Center for Transcriptional Medicine, Department of Pathology (University of Pittsburgh). The Institutional Review Board at the University of Pittsburgh has approved our protocol and given the Not Human Research Determination (IRB# STUDY24020093). Normal adult human liver cells were obtained from Human Liver Tissue & Hepatocytes Research Resource (HLTHRR) program at University of Pittsburgh. All participants provided consent and the protocol was reviewed and approved (IRB # STUDY20040276). Hepatocytes were isolated using a three-step collagenase digestion technique as previously described40. Cell viability was assessed after isolation using trypan blue exclusion, and only cell preparations with viability > 80% were used for the analysis.

### Reagents and kits

Hepatocyte-specific AAV8-TBG-Cre recombinase (AAV8-TBG-Cre; # 107787-AAV8) and the control AAV8-TBG-GFP vector (#105535-AAV8) were obtained from Addgene. GFP (catalog no. AG13105-CH) and mouse Yy1 expression plasmids were custom synthesized by Sino Biological. Wild-type Casp8 (plasmid no. 11817) and Src (plasmid no. 13663) expression plasmids were purchased from Addgene. Plasmids encoding the amino-terminal death effector domains (DEDs) of caspase-8 were generated as previously described[39]; YY1-S247A, YY1-S247D, Src-Y418D, CASP8-D73A, and DED-D73A expression constructs were generated by site-directed mutagenesis using the In-Fusion Snap Assembly Master Mix Kit (Takara, catalog no. 638945) according to the manufacturer’s instructions. All mutations were confirmed by Sanger sequencing. Primer sequences used for mutagenesis are listed in Supplementary Table 2. Plasma alanine aminotransferase (ALT) activity was measured using a commercially available assay kit (TECO Diagnostics, catalog no. A526-120) according to the manufacturer’s instructions.

### Liver primary cell isolation

Primary mouse hepatocytes were isolated from 8–10-week-old male C57BL/6J mice by collagenase perfusion[8]. Briefly, livers were perfused through the portal vein with prewarmed HBSS perfusion buffer (Corning, catalog no. 21-022-CV) supplemented with 25 mM HEPES (Sigma, catalog no. H0887) and 0.5 mM EDTA (Invitrogen, catalog no. 15575-038), followed by digestion with HBSS containing collagenase D (4.4 U per mouse; Roche, catalog no. 11088882001). The digested livers were gently dissociated, and hepatocytes were collected by low-speed centrifugation (50 × g, 1 min, 4 °C). Cells were seeded onto collagen-coated culture dishes and maintained in DMEM supplemented with 10% fetal bovine serum and 1% penicillin–streptomycin before subsequent experiments.

### Cell Culture

AML12 mouse hepatocytes (ATCC, CRL-2254) were maintained in DMEM/F12 medium (Life Technologies, catalog no. 11320) supplemented with 10% heat-inactivated fetal bovine serum (Gibco, catalog no. 16140-071) and 1× penicillin–streptomycin (Corning, catalog no. 30-002-CI). Primary mouse hepatocytes were isolated from 8–10-week-old male C57BL/6J mice as described above and cultured in DMEM containing 10% fetal bovine serum before experimental treatments. Primary human hepatocytes were maintained in hepatocyte maintenance medium (Lonza, catalog no. CC-3197) according to the manufacturer’s instructions. Following treatment, cells were harvested in Laemmli sample buffer (Bio-Rad, catalog no. 1610737) supplemented with 2-mercaptoethanol (Bio-Rad, catalog no. 161-0710) for immunoblot analysis or in RNA lysis buffer (Qiagen, catalog no. 79216) for RNA isolation and quantitative PCR.

### siRNA-mediated gene silencing and plasmid transfection

Gene silencing was performed using Dicer-substrate siRNAs (DsiRNAs; Integrated DNA Technologies). Scrambled control or gene-specific DsiRNAs were transfected into AML12 cells or primary hepatocytes using Lipofectamine RNAiMAX (Thermo Fisher Scientific) according to the manufacturer’s instructions. Briefly, cells were seeded in 24-well plates at 30–40% confluence and transfected with 40 nM DsiRNA using 1.5 μL Lipofectamine RNAiMAX in 0.5 mL culture medium. Cells were incubated for 18 h before replacement with fresh culture medium. DsiRNA sequences are listed in Supplementary Table 3. For plasmid transfection, AML12 hepatocytes were transfected using Lipofectamine LTX with PLUS Reagent (Thermo Fisher Scientific, catalog no. 15338100) according to the manufacturer’s instructions. Briefly, cells at 30–40% confluence were transfected with 0.5 μg plasmid DNA, 2 μL Lipofectamine LTX, and 0.5 μL PLUS Reagent per well of a 24-well plate. Culture medium was replaced after overnight incubation, and the cells were harvested 48 h after transfection. Primary human hepatocytes were transfected immediately after isolation while in suspension. Briefly, primary human hepatocytes were transfected in suspension by mixing DsiRNAs with the transfection reagent immediately before plating onto collagen-coated 12-well plates. Cells were cultured for 24 h before RNA or protein isolation.

### Blood and plasma analyses

Blood glucose levels were measured using a glucose meter (OneTouch Ultra, LifeScan) in mice following a 4-h fast with free access to water. Plasma alanine aminotransferase (ALT) activity was measured using the assay described in the Reagents and Kits section. Data were acquired using SoftMax Pro Software (version 5.4.1).

### Liver histology

Liver tissues were fixed in 10% neutral-buffered formalin for at least 24 h at room temperature, embedded in paraffin, and sectioned at 5 μm thickness. Sections were stained with hematoxylin and eosin (H&E) or Picrosirius Red (Polysciences, catalog no. 24901) according to the manufacturer’s instructions.

### Liver immunohistochemistry

Paraffin-embedded liver sections were deparaffinized in xylene, rehydrated through graded ethanol, and subjected to heat-induced antigen retrieval in Antigen Unmasking Solution (Vector Laboratories, catalog no. H-3300) using a pressure cooker for 10 min. Endogenous peroxidase activity was quenched with 3% hydrogen peroxide for 10 min, followed by blocking with Serum-Free Protein Block (Cell Signaling Technology, #15019) for 30 min at room temperature. Sections were incubated overnight at 4 °C with anti-COL1A1 antibody (Cell Signaling Technology, #72026; 1:500 dilution), followed by incubation with SignalStain Boost IHC Detection Reagent (Cell Signaling Technology, #8114). Immunoreactivity was visualized using the DAB substrate kit (Cell Signaling Technology, #8059). Sections were counterstained with hematoxylin, dehydrated, mounted, and imaged using a Nikon bright-field microscope.

### Immunofluorescence microscopy

AML12 hepatocytes or primary hepatocytes were fixed in 4% paraformaldehyde for 10 min at room temperature, washed with phosphate-buffered saline (PBS), and incubated overnight at 4 °C with primary antibodies against HA (Cell Signaling Technology, #3724; 1:500 dilution) or YY1 (Cell Signaling Technology, #46395; 1:1,000 dilution). Cells were then washed, incubated with Alexa Fluor-conjugated secondary antibodies and DAPI, and imaged in PBS using an Olympus DP74 fluorescence microscope. For liver tissue, paraffin-embedded sections were deparaffinized, rehydrated, and subjected to heat-induced antigen retrieval in Antigen Unmasking Solution (Vector Laboratories, #H-3300) using a pressure cooker for 10 min. After blocking with Immunofluorescence Blocking Buffer (Cell Signaling Technology, #12411), sections were incubated overnight at 4 °C with antibodies against α-smooth muscle actin (α-SMA; Abclonal, #A7248; 1:300 dilution), F4/80 (Cell Signaling Technology, #70076; 1:500 dilution), or YY1 (Cell Signaling Technology, #46395; 1:500 dilution). Sections were then incubated with Alexa Fluor 555-conjugated donkey anti-rabbit secondary antibody (Invitrogen, #A-31572; 1:500 dilution), mounted using DAPI-containing mounting medium (Thermo Fisher Scientific, #P36935), and imaged using an Olympus DP74 fluorescence microscope with cellSens software.

### Image analysis

Histological and immunofluorescence images were analyzed using ImageJ2 (version 2.14.0). Identical threshold settings were applied to all images within each experiment. For liver sections, six randomly selected fields were analyzed per section, and the mean value for each mouse was used for statistical analysis. Nuclear YY1 fluorescence intensity was quantified from DAPI-positive nuclei using the same analysis parameters across all groups.

### Immunoblotting

Liver tissues were homogenized in RIPA buffer (Thermo Fisher Scientific, #89900) supplemented with protease and phosphatase inhibitors. Protein concentrations were determined using the bicinchoninic acid (BCA) assay (Thermo Fisher Scientific, #23227). Equal amounts of protein were separated on 4–20% Tris–glycine gels (Invitrogen, #EC60285) and transferred onto nitrocellulose membranes (Bio-Rad, #1620115). Membranes were blocked for 30 min at room temperature in Tris-buffered saline containing 0.1% Tween-20 (TBST) and 5% non-fat milk, followed by incubation overnight at 4 °C with primary antibodies diluted 1:1,000 in blocking buffer. Primary antibodies included anti-YY1 (Cell Signaling Technology, #46395), anti-phosphoserine (MilliporeSigma, #AB1603), anti-phospho-Src (Tyr416; Cell Signaling Technology, #59548), anti-Src (Cell Signaling Technology, #2109), anti-phospho-GSK3β (Tyr216; Proteintech, #29125), anti-GSK3β (Cell Signaling Technology, #12456), anti-caspase-8 (Cell Signaling Technology, #4790), anti-HA (Cell Signaling Technology, #3724), anti-β-actin (Cell Signaling Technology, #5125), and anti-GAPDH (Cell Signaling Technology, #3683). Immunoreactive bands were detected using horseradish peroxidase-conjugated donkey anti-rabbit IgG secondary antibody (Jackson ImmunoResearch, #711-035-152; 1:5,000 dilution) and SuperSignal West Pico chemiluminescent substrate (Thermo Fisher Scientific, #34080). For cultured cells, proteins were extracted directly into Laemmli sample buffer (Bio-Rad, #161-0737) containing 5% 2-mercaptoethanol, heated at 100 °C for 5 min, and subjected to SDS–PAGE and immunoblotting as described above.

### Immunoprecipitation assays

Cells were lysed in ice-cold Cell Lysis Buffer (Cell Signaling Technology, #9803) supplemented with protease and phosphatase inhibitor cocktails (Cell Signaling Technology, #5872). Lysates were clarified by centrifugation at 14,000 *g* for 10 min at 4 °C, and protein concentrations were determined using the BCA assay. Equal amounts of protein (approximately 1 mg) were incubated overnight at 4 °C with anti-Src (Cell Signaling Technology, #2109; 1:100 dilution) or anti-HA (Cell Signaling Technology, #3724; 1:100 dilution) antibodies. Immune complexes were captured using Protein A magnetic beads (Cell Signaling Technology, #73778) for 2 h at 4 °C, washed five times with ice-cold lysis buffer, and eluted by boiling in SDS sample buffer for 5 min. Eluted proteins were resolved by SDS–PAGE and analyzed by immunoblotting as described above, except that TrueBlot secondary antibodies (eBioscience) were used for detection. Where indicated, nuclear proteins were isolated using the Cell Fractionation Kit (Cell Signaling Technology, #9038) before immunoprecipitation. For endogenous YY1 immunoprecipitation, lysates were incubated overnight with anti-YY1 antibody (Cell Signaling Technology, #46395; 1:100 dilution) before capture with Protein A magnetic beads.

### Quantitative RT-qPCR

Total RNA was isolated from liver tissue or cultured hepatocytes using the RNeasy Mini Kit (Qiagen, #74106) according to the manufacturer’s instructions. RNA concentration was determined using a NanoDrop spectrophotometer (Thermo Fisher Scientific). First-strand cDNA was synthesized from 1 μg of total RNA using SuperScript II Reverse Transcriptase (Invitrogen, #4368814) with oligo(dT) primers. Quantitative PCR was performed on a QuantStudio 3 Real-Time PCR System (Applied Biosystems) using SYBR Green Master Mix (Thermo Fisher Scientific, #4367659). Relative gene expression was calculated using the comparative Ct (2^−ΔΔCt^) method after normalization to *Hprt* expression. Primer sequences are provided in Supplementary Table 3.

### LC–MS/MS identification of YY1 Ser247 phosphorylation

Immunoprecipitated samples were separated on 4–12% gradient SDS-PAGE gels and stained with SimplyBlue (Thermo Fisher Scientific). Protein bands were excised, and in-gel digestion was conducted as described previously[40], with minor modifications. Gel slices were washed with 1:1 (acetonitrile: 100 mM ammonium bicarbonate) for 30 min, dehydrated with 100% acetonitrile for 10 min until shrunken, and dried in a SpeedVac for 10 min without heat. Reduction was performed with 5 mM DTT for 30 min at 56 °C in an air thermostat, followed by cooling to room temperature. Alkylation was carried out with 11 mM IAA for 30 min in the dark. Gel slices were sequentially washed with 100 mM ammonium bicarbonate and 100% acetonitrile for 10 min each, dried in a SpeedVac, and rehydrated in 25 ng/μL trypsin in 50 mM ammonium bicarbonate on ice for 30 min. Digestion was performed overnight at 37 °C in an air thermostat. Peptides were extracted from the gel slices with extraction buffer (1:2, v/v, 5% formic acid/acetonitrile) under high-speed shaking.

Supernatants from the digestion and extraction steps were combined and dried in a SpeedVac. Peptides were resuspended in 3% acetonitrile/0.1% formic acid before LC–MS/MS analysis. Peptides were separated over a 40-min gradient at a flow rate of 400 nl/min on a reverse-phase C18 column with an integrated CaptiveSpray emitter (25 cm × 75 µm, 1.6 µm; IonOpticks). Mobile phase A consisted of 0.1% formic acid in water, and mobile phase B consisted of 0.1% formic acid in acetonitrile. The gradient was programmed as follows: 2% to 25% B over 30 min, 25% to 40% B over 5 min, then increased to 95% B before column re-equilibration. The timsTOF Pro2 was operated in PASEF mode (2) with the following settings: Mass Range 100 to 1700 m/z, 1/K0 range 0.6 to 1.6, Ramp time 100 ms, Lock Duty Cycle to 100%, Capillary Voltage 1600 V, Dry Gas 3 l/min, Dry Temp 200°C, PASEF settings: 10 MS/MS Frames (1.16 seconds duty cycle), charge range 0-5, the active exclusion for 0.4 min, Target intensity 20000, Intensity threshold 2500, CID collision energy 59 eV. A polygon filter was applied to the m/z and ion mobility plane to select features most likely representing peptide precursors rather than singly charged background ions. Acquired PASEF raw files were analyzed using MaxQuant (version 2.6.5.0) with the integrated Andromeda search engine using the default search parameters[41]. The first search peptide mass tolerance was set to 20 ppm, and the main search peptide mass tolerance was set to 4.5 ppm. MS/MS spectra were searched against the Mus musculus reference proteome (UniProt proteome ID: UP000000589). Enzyme specificity was set to trypsin/P, allowing up to two missed cleavages. Carbamidomethylation of cysteine residues (+57.021 Da) was specified as a fixed modification, whereas oxidation of methionine (+15.995 Da), phosphorylation of serine, threonine, and tyrosine (STY), and deamidation of asparagine and glutamine (+0.984 Da) were included as variable modifications. The false discovery rate (FDR) was controlled at 1% at the peptide-spectrum match (PSM), peptide, and protein levels using a target-decoy approach. A minimum of one unique peptide was required for protein identification. Protein intensity data generated by MaxQuant were used for downstream statistical analysis.

### Quantification and statistical analysis

Statistical analyses were carried out using GraphPad Prism (version 9.4). Data were tested for normality using the Shapiro-Wilk test. Data that passed the normality test were analyzed using the two-tailed Student’s t-test for comparison of two groups or one- or two-way analysis of variance (ANOVA) with Bonferroni post hoc analysis for comparison of more than two groups. Data are shown as mean ± SEM and differences were considered statistically significant at p < 0.05. No statistical methods were used to pre-determine sample sizes, but our sample sizes are similar to those reported in previous publications[9, 19, 38]. No animals or data points were excluded from the analyses. The investigators conducting the experiments were not blinded to the identity of the experimental groups.

## Data availability

All data associated with this study are present in the paper or the Supplementary Materials. The raw LC-MS/MS data have been deposited in the PRIDE database under accession number PXD082147. All the databases and datasets used in the study along with appropriately accessible links in the manuscript. All data supporting the figures in this paper are provided in the source data published with the article.

## Code availability

No original code was created for this study.

## Author contributions

X.W. and I.T. conceived the study and wrote the manuscript, with all other authors providing comments and editing. X.W. and E.J.J. conducted the experiments. L.A.P.F. and A.S.G gave advice and reagents for the human hepatocyte experiments. D.G.S. was instrumental for the studies focused on the DED domains of caspase-8.

## Supplemental Information

**Supplementary Fig. 1.**
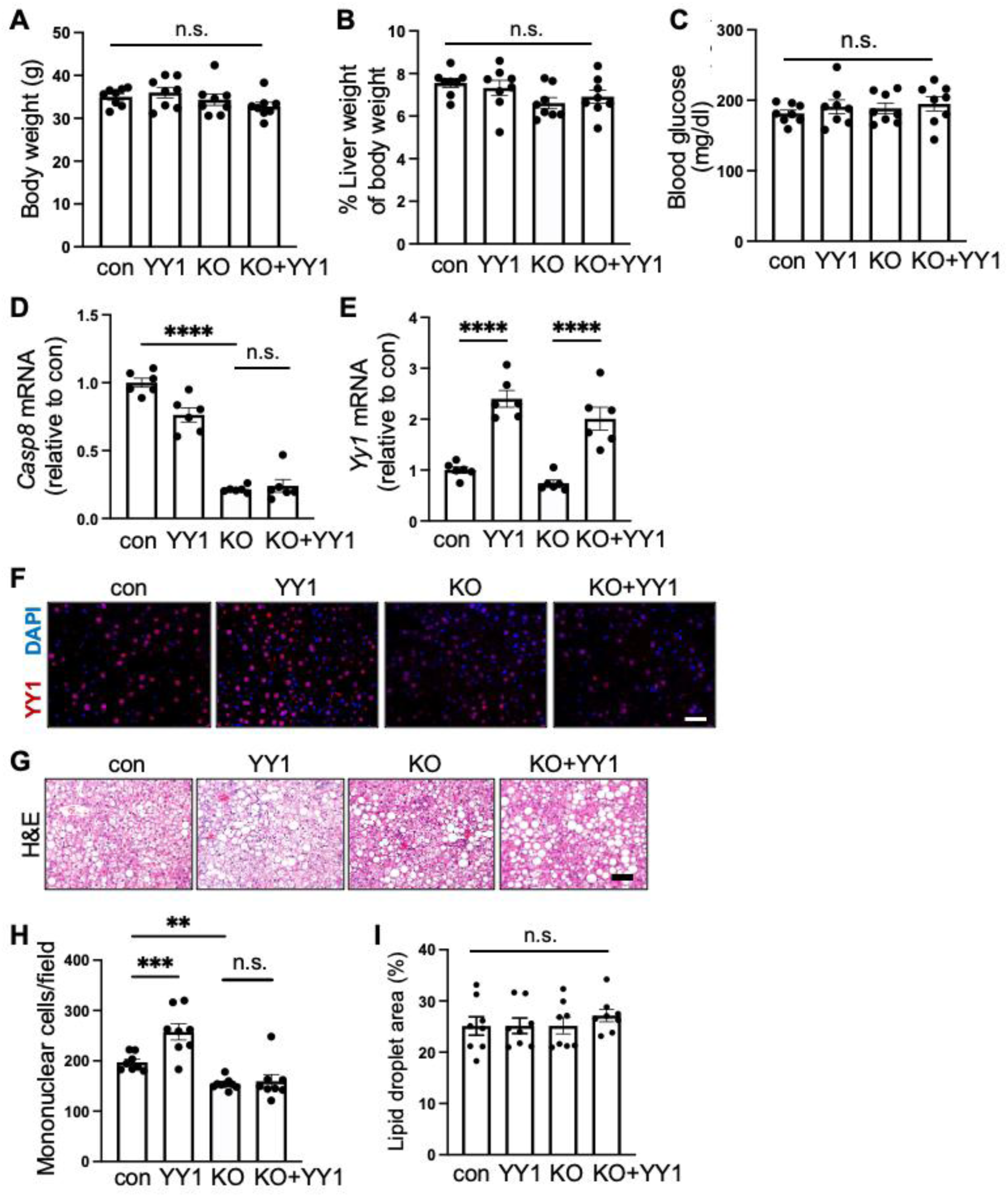
Additional characterization of YY1 overexpression in wild-type and hepatocyte-caspase-8 knockout MASH mice. Related to Fig. 1. *Casp^fl/fl^* mice were fed the FPC diet for 10 weeks and then injected with AAV8-TBG-Cre to generate hepatocyte-specific caspase-8 knockout mice (H-C8-KO) or co-injected with AAV8-TBG-Yyl to overexpress YY1 in hepatocytes (H-C8-KO+YY1). Mice remained on the FPC diet for an additional 6 weeks before analysis. **A-C,** Body weight (A), liver weight as a percentage of body weight (B), and blood glucose levels measured 4 h after food withdrawal (C) in the four groups of mice shown in Fig. 1. **D-E,** mRNA expression of *Casp8* (D) and *Yyl* (E). **F**, YY1 immunofluorescence images (red), with DAPI nuclear counterstain (blue); bar, 100 pm. **G,** H&E staining of liver sections; bar, 200 pm. **H,** Quantification of hepatic mononuclear cells. **I,** Quantification of lipid droplet area. Data are presented as mean ± s.e.m. Statistical analysis was performed using one-way ANOVA; **A-C, H-I,** n = 8 mice per group, **D-E,** n = 6 mice per group. **p<0.01, ***p<0.001, ****p<0.0001, n.s., not significant.

**Supplementary Fig. 2.**
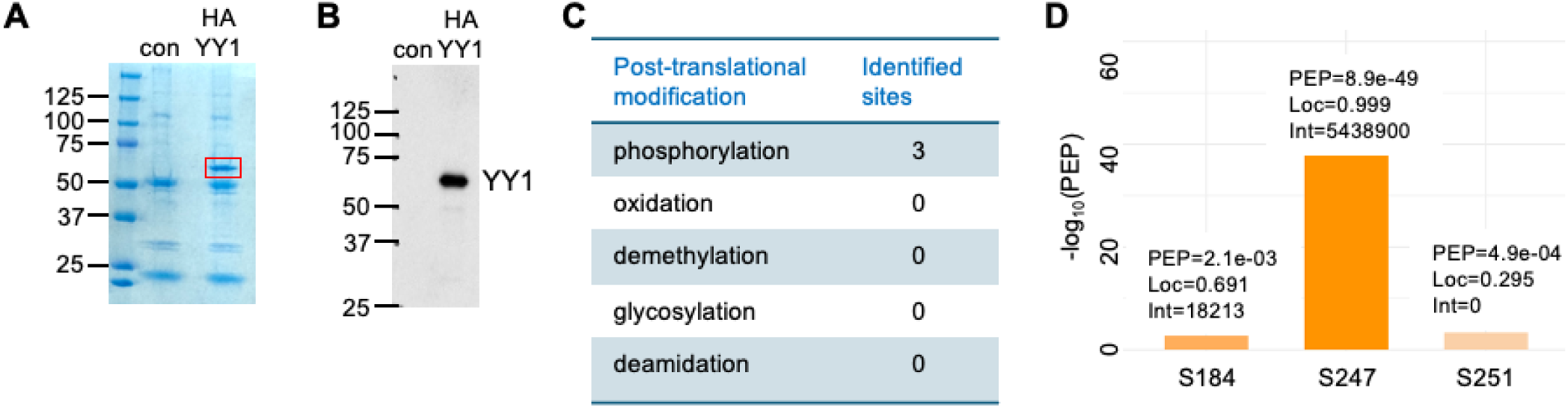
Identification of YY1 phosphorylation sites by LC-MS/MS. Related to Fig. 2. **A,** YY1-knockout AML12 hepatocytes were transfected with GFP (control) or HA-tagged YY1. Nuclear extracts were prepared, HA-YY1 was immunoprecipitated (IP), and the IP products were separated by SDS-PAGE followed by Coomassie Blue staining. **B,** Immunoblot analysis of the HA immunoprecipitates using an anti-YYl antibody. **C,** Summary of YY1 post-translational modifications identified by LC–MS/MS analysis. **D,** LC-MS/MS analysis identified three candidate phosphorylation sites on YY1. Candidate sites were ranked according to posterior error probability (PEP), localization probability (Loc), and peptide intensity (Int), with Ser247 identified as the highest-confidence phosphorylation site.

**Supplementary Fig. 3.**
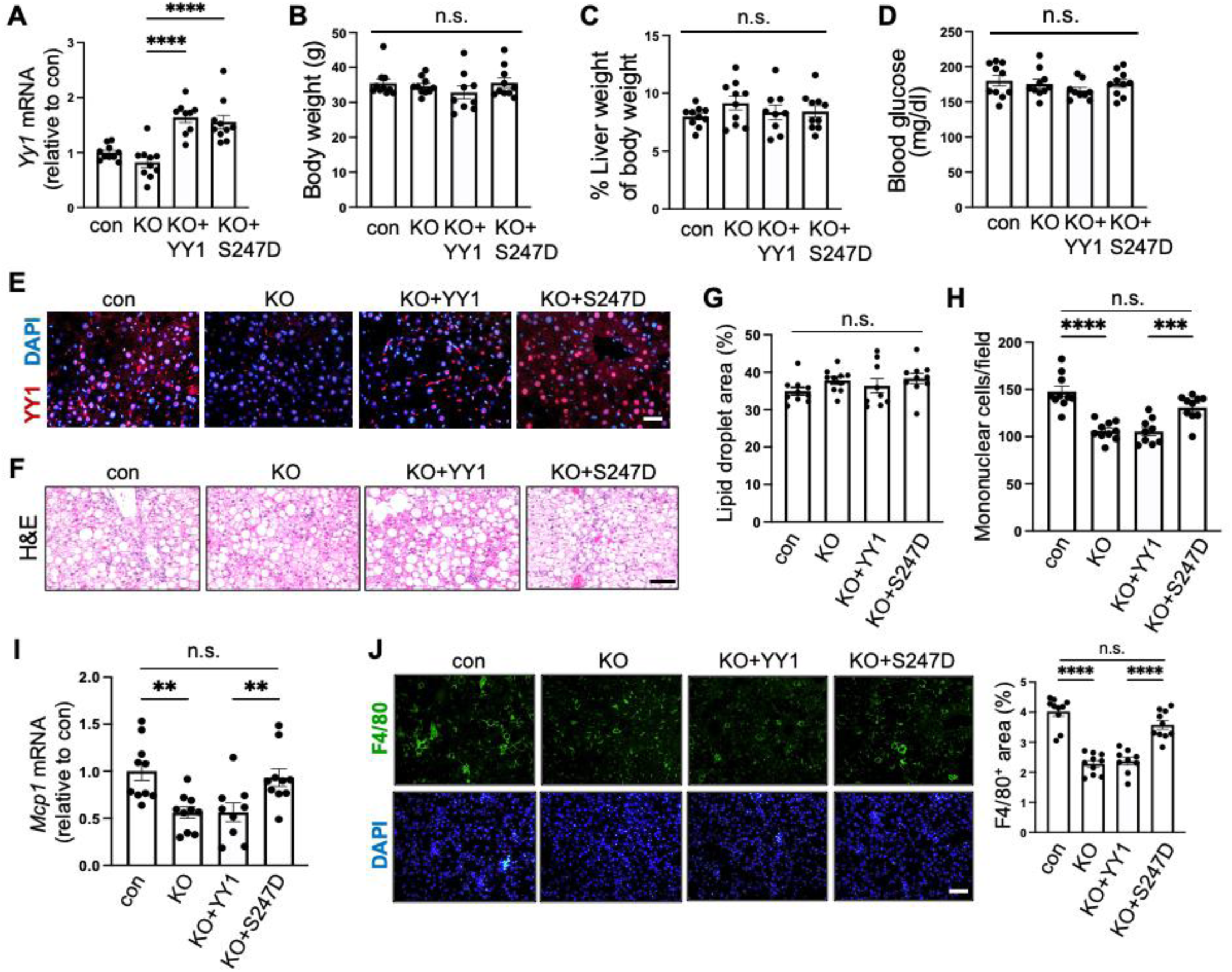
Additional characterization of wild-type and constitutively active YY1 overexpression in hepatocyte-caspase-8 knockout MASH mice. Related to Fig. 6. *Casp^fl/fl^ mice* were fed the FPC diet for 10 weeks, followed by hepatocyte-specific deletion of Casp8 using AAV8-TBG-Cre injection (KO). Mice were subsequently co-injected with AAV8-TBG vectors expressing either wild-type YY1 (KO+YY1) or constitutively active YY1 (KO+S247D) and maintained on the FPC diet for an additional 6 weeks before analysis. **A,** Hepatic mRNA expression of *Yyl.* **B,** Body weight. **C,** Liver weight as a percentage of body weight. **D,** Blood glucose levels measured 4 h after food withdrawal. **E,** YY1 immunofluorescence (red), with nuclear DAPI counterstain; bar, 100 pm. **F,** H&E staining of liver sections; bar, 200 pm, **G,** Quantification of lipid droplet area. **H,** Quantification hepatic mononuclear cells. **I,** mRNA expression of *Mcpl.* **J,** F4/80 immunofluorescence (red; arrows), with nuclear DAPI counterstain (blue), and quantification of F4/80 positive area; bar, 200 pm. Data are presented as mean ± s.e.m. Statistical significance was determined by one-way ANOVA. Sample sizes: control, n = 10; KO, n = 10; KO + YY1, n = 9; KO + S247D, n = 10. **p < 0.01, ***p < 0.001, ***p < 0.0001; n.s., not significant.

**Supplementary Fig. 4.**
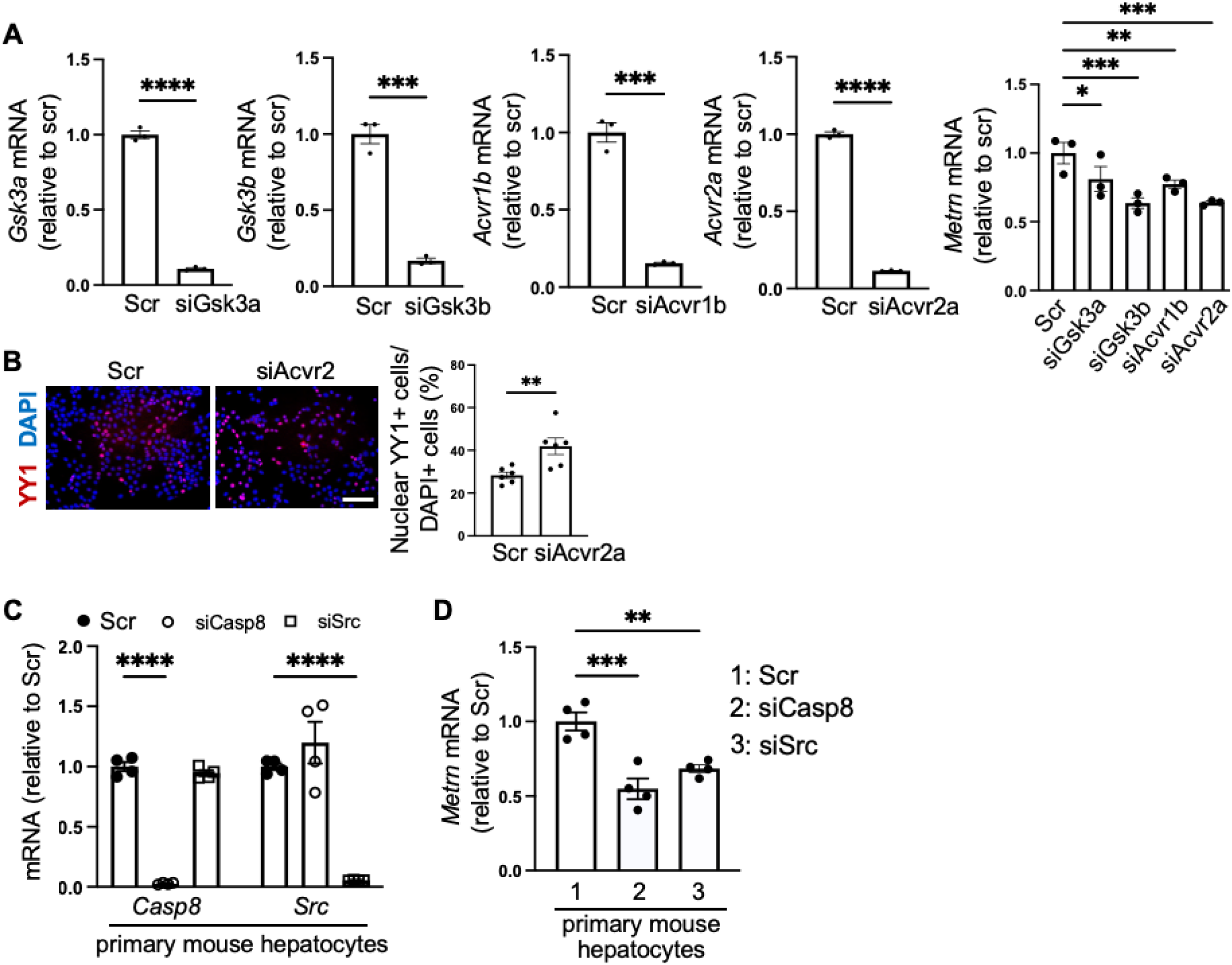
Identification of GSK3β as the upstream regulator of YY1 in AML12 hepatocytes and primary mouse hepatocytes. Related to Fig.4. **A,** *Gsk3a, Gsk3b, Acvrlb, Acvr2a,* and *Metm* mRNA expression in AML 12 hepatocytes transfected with scrambled (Scr) RNA or siRNAs targeting *Gsk3a, Gsk3b, Acvrlb,* or *Acvr2a.* n = 3 biologically independent experiments per group. Statistical significance was determined by two-tailed unpaired Student’s t-test and one-way ANOVA (right). **B,** Immunofluorescence microscopy images and quantification of YY1-positive nuclei in AML12 hepatocytes treated with scrambled RNA (Scr) or siAcvr2a. Scale bar, 100 pm. n = 6 biologically independent experiments per group. Statistical significance was determined by two-tailed unpaired Student’s t-test. **C,** *Casp8* and *Src* mRNA expression in primary mouse hepatocytes treated with scrambled RNA (Scr), siCasp8, or siSrc. n = 4 biologically independent experiments per group. Statistical significance was determined by one-way ANOVA. **D,** *Metrn* mRNA expression in the cells shown in panel c. n = 4 biologically independent experiments per group. Statistical significance was determined by one-way ANOVA. Data are presented as mean ± s.e.m. *p<0.05, **p<0.01, ***p<0.001, ****p<0.0001.

**Supplementary Fig. 5.**
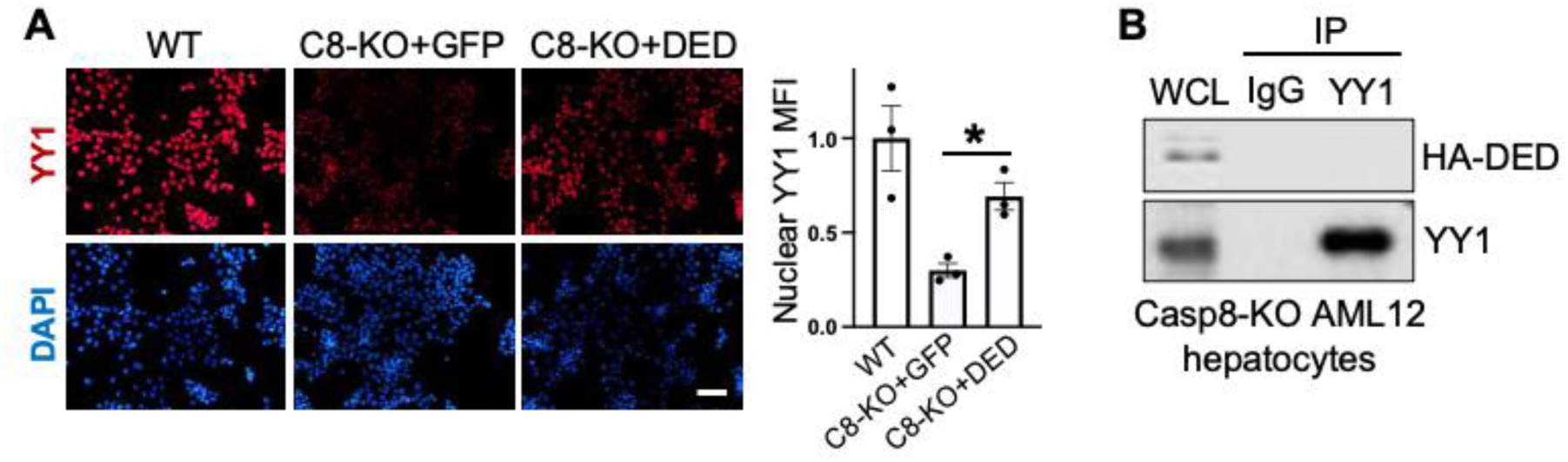
Additional evidence that the caspase-8 DED domain activates the Src-GSK3β-YYl signaling pathway. Related to Fig, 5. **A,** Immunofluorescence microscopy images and relative MFI quantification of nuclear YY1 in wild-type (WT) or caspase-8-KO (C8-KO) AML12 hepatocytes transfected with control GFP or DED. Scale bar, 200 μm. n = 3 biologically independent experiments per group. **B,** YY1 was immunoprecipitated from HA-DED-transfected caspase-8-KO AML12 hepatocytes and immunoblotted for YY1 and HA in the immunoprecipitates (IP) and whole-cell lysates (WCL). Statistical significance was determined by one-way ANOVA. Data are presented as mean ± s.e.m. *p<0.05.

**Supplementary Table 1.** (related to Figs. 5f, 5g, 5h, 5i). Pathology of human MASH liver samples.

| ID | Age | Gender | Pathological diagnoses |
| --- | --- | --- | --- |
| 1 | 68 | Male | Non-alcoholic steatohepatitis; cirrhosis, mixed macronodular and micronodular type. Minimal inflammatory activity, with minimal or no ongoing steatosis. No evidence of hepatocyte dysplasia or malignancy. |
| 2 | 56 | Female | Type 2 Diabetes. Obesity. Advanced chronic liver disease (stage 4: "cirrhosis"). Marked fibrosis throughout the entire liver. The residual hepatocellular parenchyma shows abundant well-formed Mallory hyaline, consistent with advanced chronic liver disease from steatohepatitis. |
| 3 | 53 | Male | MASH. Cirrhosis. Mild steatosis and focal active steatohepatitis. Focal Mallory hyaline is identified. |
| 4 | 67 | Female | Cirrhosis, MASH. Active steatohepatitis, grade 2 of 3. |
| 5 | 62 | Female | Diseased (MASH) – Child-Pugh-Score: B (scoring:7)<br>Serum albumin (g/dL):3.7; total Bilirubin (mg/dL) :1.0; International Normalized Ratio (INR):1.0<br>Ascites (yes) and Encephalopathy (yes)<br>METAVIR Score = F4 |
| 6 | 55 | Female | Diseased (MASH) – Child-Pugh-Score: C (scoring:10)<br>Serum albumin (g/dL):2.6; total Bilirubin (mg/dL):0.6; International Normalized Ratio (INR):1.1<br>Ascites (yes) and Encephalopathy (yes)<br>METAVIR Score = F4 |
| 7 | 53 | Female | Diseased (MASH) – Child-Pugh-Score: B (scoring: 9)<br>Serum albumin (g/dL):3.2; total Bilirubin (mg/dL):2.9; International Normalized Ratio (INR):1.3<br>Ascites (no) and Encephalopathy (yes)<br>METAVIR Score = F4 |

Sample 1-4: MASH; 5-7: MASH hepatocyte isolation.

**Supplementary Table 2.** (related to multiple figures). DsiRNA and crRNA sequences used in this study.

| **Target Gene** | **Oligo information** | **Sequences (5' to 3')** |
| --- | --- | --- |
| *Casp8* | mouse DsiRNA | rGrGrArArCrCrUrGrGrUrArUrArUrUrCrArGrUrCrArCrUTT |
| *CASP8* | human DsiRNA | rGrUrCrArUrGrCrUrCrUrArUrCrArGrArUrUrUrCrArGrAAG |
| *Src* | mouse DsiRNA | rUrUrUrArCrGrUrCrCrArCrUrGrArGrCrCrArCrArGrUrACC |
| *SRC* | human DsiRNA | rGrCrArArGrGrUrGrCrCrArArArUrUrCrCrCrCrArUrCrAAG |
| *Gsk3b* | mouse DsiRNA | rCrArGrArUrCrArUrGrArGrArArArGrCrUrArGrArCrCrACT |
| *Gsk3a* | mouse DsiRNA | rCrArArGrArArGrUrGrGrCrUrUrArCrArCrUrGrArCrArUCA |
| *Acvr1b* | mouse DsiRNA | rCrGrArGrArCrArArUrCrArArCrArUrGrArArGrCrArCrUTT |
| *Acvr2a* | mouse DsiRNA | rGrCrArUrUrUrCrUrGrArGrGrArUrArGrArUrArUrGrUrACG |
| *Casp8* | mouse crRNA | /AltR1/rCrUrUrCrCrUrArGrArCrUrGrCrArArCrCrGrArGrGrUrUrUrUrArGrArGrCrUrA<br>rUrGrCrU/AltR2/ |
| *Yy1* | mouse crRNA | /AltR1/rGrCrCrCrArCrCrArCrCrGrUrGrGrUrCrUrCrGrArGrUrUrUrUrArGrArGrCrUrA<br>rUrGrCrU/AltR2/ |

**Supplementary Table 3.**
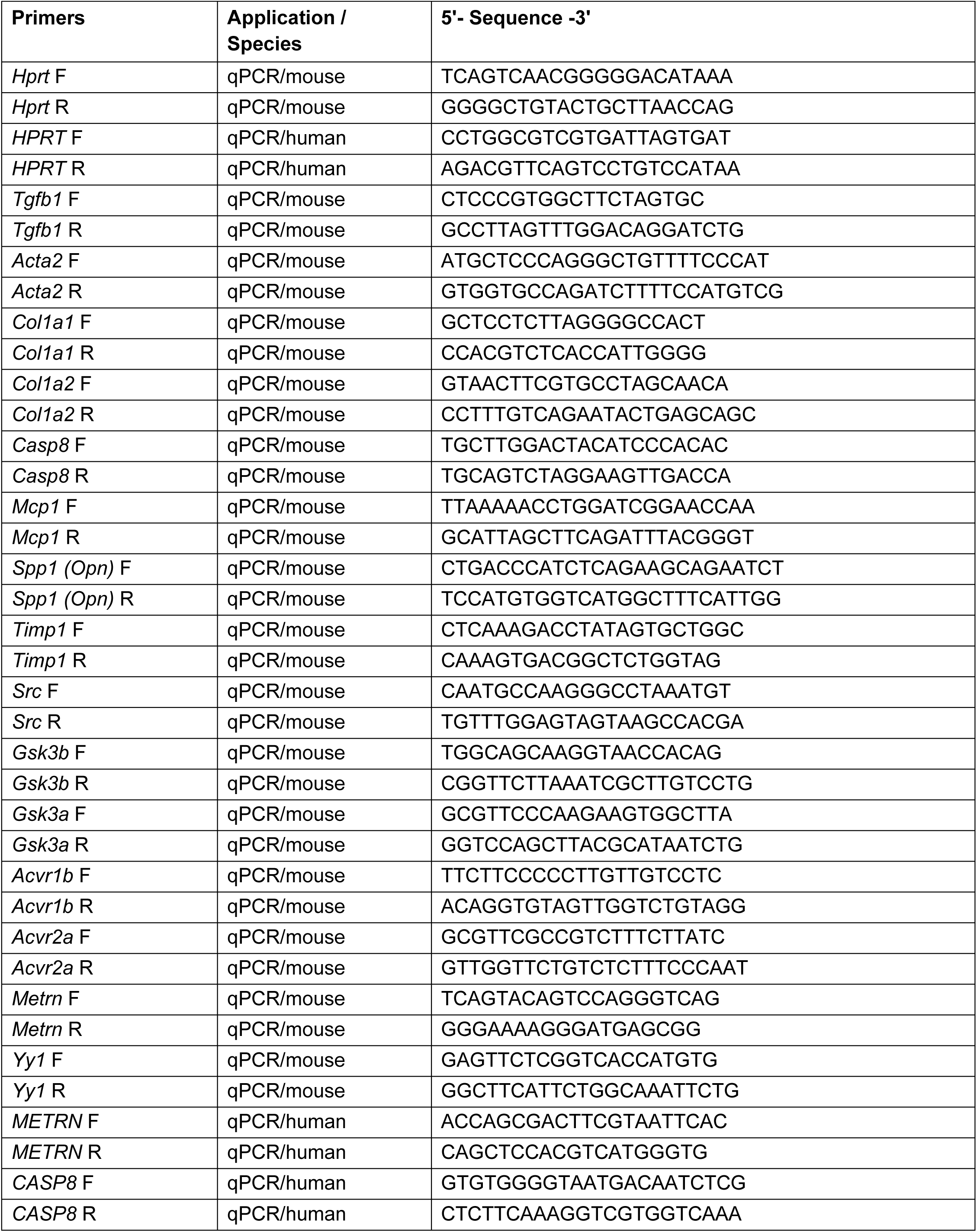
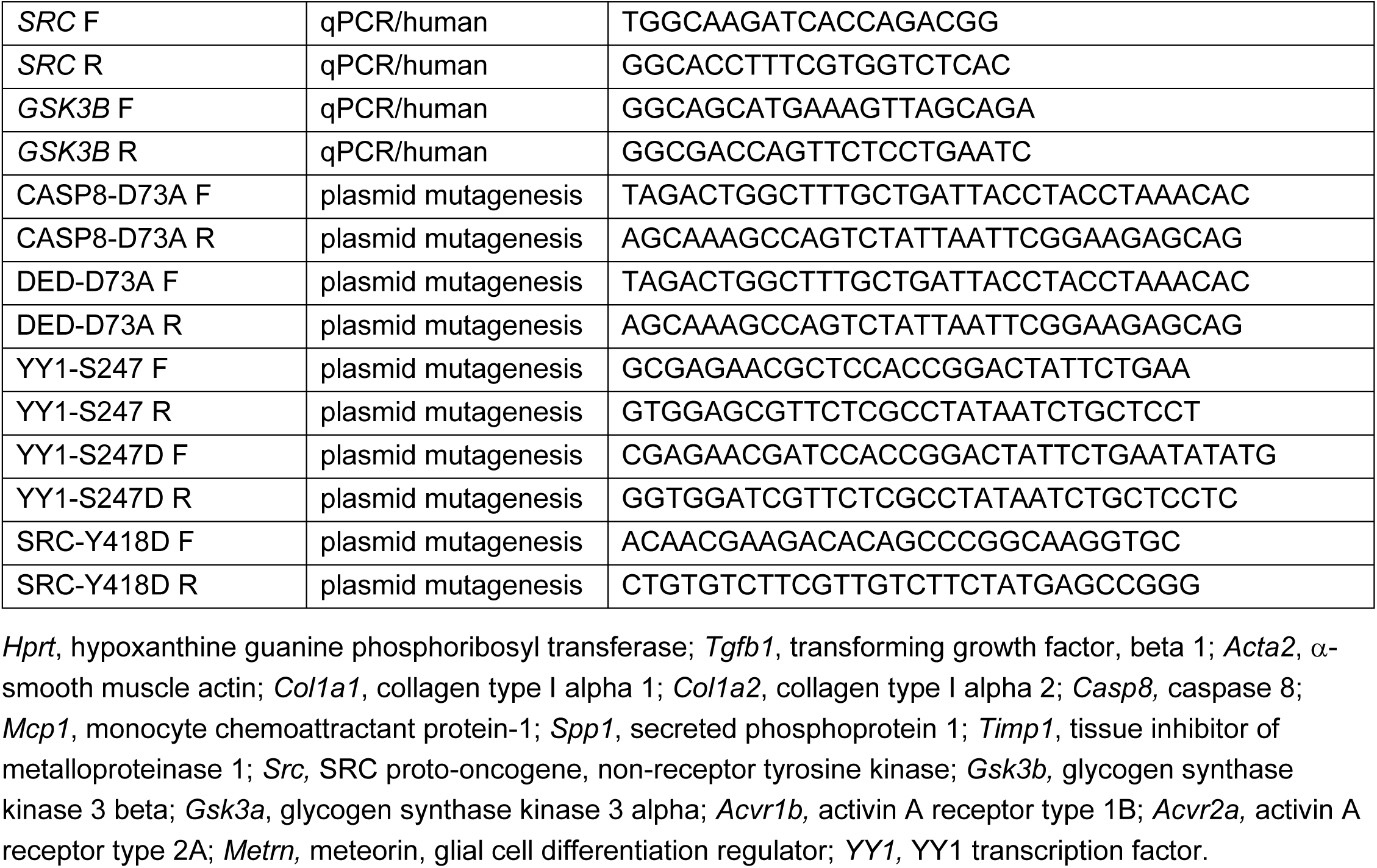
(Related to multiple figures). Primer sequences used for quantitative real-time PCR (RT-qPCR), PCR, and site-directed mutagenesis.

